# Prion protein lowering is a disease-modifying therapy across prion disease stages, strains, and endpoints

**DOI:** 10.1101/2020.03.27.011940

**Authors:** Eric Vallabh Minikel, Hien T Zhao, Jason Le, Jill O’Moore, Rose Pitstick, Samantha Graffam, George A Carlson, Michael P Kavanaugh, Jasna Kriz, Jae Beom Kim, Jiyan Ma, Holger Wille, Judd Aiken, Deborah McKenzie, Katsumi Doh-ura, Matthew Beck, Rhonda O’Keefe, Jacquelyn Stathopoulos, Tyler Caron, Stuart L Schreiber, Jeffrey B Carroll, Holly B Kordasiewicz, Deborah E Cabin, Sonia M Vallabh

## Abstract

Lowering of prion protein (PrP) expression in the brain is a genetically validated therapeutic hypothesis in prion disease. We recently showed that antisense oligonucleotide (ASO)-mediated PrP suppression extends survival and delays disease onset in intracerebrally prion-infected mice in both prophylactic and delayed dosing paradigms. Here, we examine the efficacy of this therapeutic approach across diverse paradigms, varying the dose and dosing regimen, prion strain, treatment timepoint, and examining symptomatic, survival, and biomarker readouts. We recapitulate our previous findings with additional PrP-targeting ASOs, and demonstrate therapeutic benefit against four additional prion strains. We demonstrate that less than 25% PrP suppression is sufficient to extend survival and delay symptoms in a prophylactic paradigm. Rise in both neuroinflammation and neuronal injury markers can be reversed by a single dose of PrP-lowering ASO administered after the detection of pathological change. Chronic ASO-mediated suppression of PrP beginning at any time up to early signs of neuropathology confers benefit similar to constitutive heterozygous PrP knockout. Remarkably, even after emergence of frank symptoms including weight loss, a single treatment prolongs survival by months in a subset of animals. These results support ASO-mediated PrP lowering, and PrP-lowering therapeutics in general, as a promising path forward against prion disease.

## Introduction

Prion disease, a rapidly fatal and currently untreatable neurodegenerative disease, is caused by the post-translational conformational corruption of host-encoded prion protein (PrP) (1). Due to its central role in disease pathophysiology, reduction of native PrP is an attractive therapeutic hypothesis in prion disease (2). Homozygous deletion of PrP prevents prion infection (3, 4), while heterozygous PrP knockout delays development of disease following prion infection (4–7) and transgenic PrP overexpression accelerates it (8), providing genetic evidence of a continuous dose-response relationship between PrP dosage and disease susceptibility. Conditional knockout systems have confirmed that post-natal depletion confers significant survival benefit, even in the presence of low levels of residual PrP expression (9, 10). Knockout animals are healthy (11–13). The only established knockout phenotype is a peripheral neuropathy, apparently due to deficiency of myelin maintenance signaling to a Schwann cell receptor (14), which is histologically evident yet phenotypically mild to undetectable in homozygotes and is not observed in heterozygotes (15, 16). Heterozygous inactivating mutations also appear to be tolerated in humans (17, 18), minimizing any concern about on-target toxicity of pharmacologic PrP lowering.

The use of therapeutic oligonucleotides to lower PrP by targeting its RNA has been considered for over two decades (19), but early attempts, hampered by drug delivery and distribution challenges, yielded modest or no benefit in animal models (20–24). Genetically targeted therapies designed to reduce levels of other single target proteins have recently shown promising target engagement in the human central nervous system (25–27). Building on these successes, we and others recently showed that PrP-lowering antisense oligonucleotides (ASOs), bolus dosed into cerebrospinal fluid (CSF), can extend survival by 61-98% in prion-infected mice (28).

For PrP-lowering therapy to advance effectively, a number of fundamental questions must be addressed. While heterozygous knockout animals show a clear benefit to 50% PrP reduction (4–7), the minimal threshold of PrP knockdown needed to confer benefit has not been established. The existence of different prions strains, or subtypes, has complicated previous drug development efforts: antiprion compounds with non-PrP-lowering mechanisms of action have failed to generalize across strains (29–33), and prions have been shown capable of adapting to drug treatment, giving rise to new drug-resistant strains (30, 34, 35). It is therefore critical to test any potential prion disease therapeutic strategy against multiple prion strains, and to monitor for development of drug-resistant prions. While our previous experiments showed the delay of pathological changes to brain tissue of ASO-treated animals (28), we did not investigate potential impact on established neuropathological changes following treatment. Further, our prior experiments relied on a limited number of ASO doses, rather than chronic dosing aiming for continuous suppression, though the latter paradigm better mirrors clinical use of ASOs. Finally, in prion disease it is important to understand at what disease stage treatment can be effective. Clinically, most prion disease patients die within half a year of first symptoms (36), and this rapid decline is mirrored by high levels of biofluid neuronal injury and prion seeding biomarkers in the symptomatic phase of disease (37–42). Meanwhile, individuals at risk for genetic prion disease, caused by protein-altering variants in the prion protein gene (*PRNP*), can be identified through predictive genetic testing when disease onset is on expectation years or decades away (43), ahead of molecular markers of pathology (44). This spectrum motivates investigation of a range of treatment timepoints relative to prion inoculation, development of molecular pathology, and presentation of frank symptoms to explore the potential of PrP-lowering treatment.

Here, using ASOs as tool compounds, we test the efficacy of PrP lowering via an RNAse-H dependent mechanism across a variety of therapeutic paradigms in prion-infected mice, in order to fill these critical knowledge gaps and inform the clinical development of PrP-lowering drugs.

## Materials and methods

### Study design

At the Broad Institute, procedures (prion infection and ASO administration) were performed by investigators (SV and EVM) with full knowledge of study design, while all behavioral observations, weights, nest scores, and final endpoint determinations were taken by veterinary technicians (primarily JL and SG, with others on an on-call basis) blinded to the animals’ treatment status or genotype. At the McLaughlin Research Institute, raters were not blinded. Disease endpoints (see below) were pre-specified at the time of protocol approval.

### ASO discovery

ASOs 1 and 2 were prioritized through a cellular screen of roughly 500 ASOs in HEPA1-6 cells, then further characterized in cells and *in vivo* as previously described (28). Briefly, ASOs were incubated with cells at 7 uM for 24 hours. RNA was then purified from harvested cells, and mouse *Prnp* mRNA was quantified using RT-PCR (Figure 1A). Potent ASOs were next subjected to a 4-point dose response experiment. Finally, C57BL/6N mice received bolus doses of active ASOs 1 and 2 to characterize potency *in vivo* (Figure 1), (28). To generate ASOs featuring a 10-base deoxynucleotide gap symmetrically flanked with 2’O-methoxyethyl (MOE) modified nucleotides, we performed optimization around the ASO 1 and 2 active sites (Figure 1B). Groups of N=4 C57BL/6N mice subsequently received a 700 μg dose of one of five new candidate ASOs, delivered by single bolus intracerebroventricular (ICV) injection. Eight weeks later, *Prnp* mRNA suppression was quantified by qPCR in cortex and thoracic cord (Figure 1C-D). Combined with animal weight and neurological exam data, collected weekly (Figure 1E), these data led to prioritization of ASOs 5 and 6. Chemical modifications for all ASOs are shown in Table 1.

**Figure 1.**
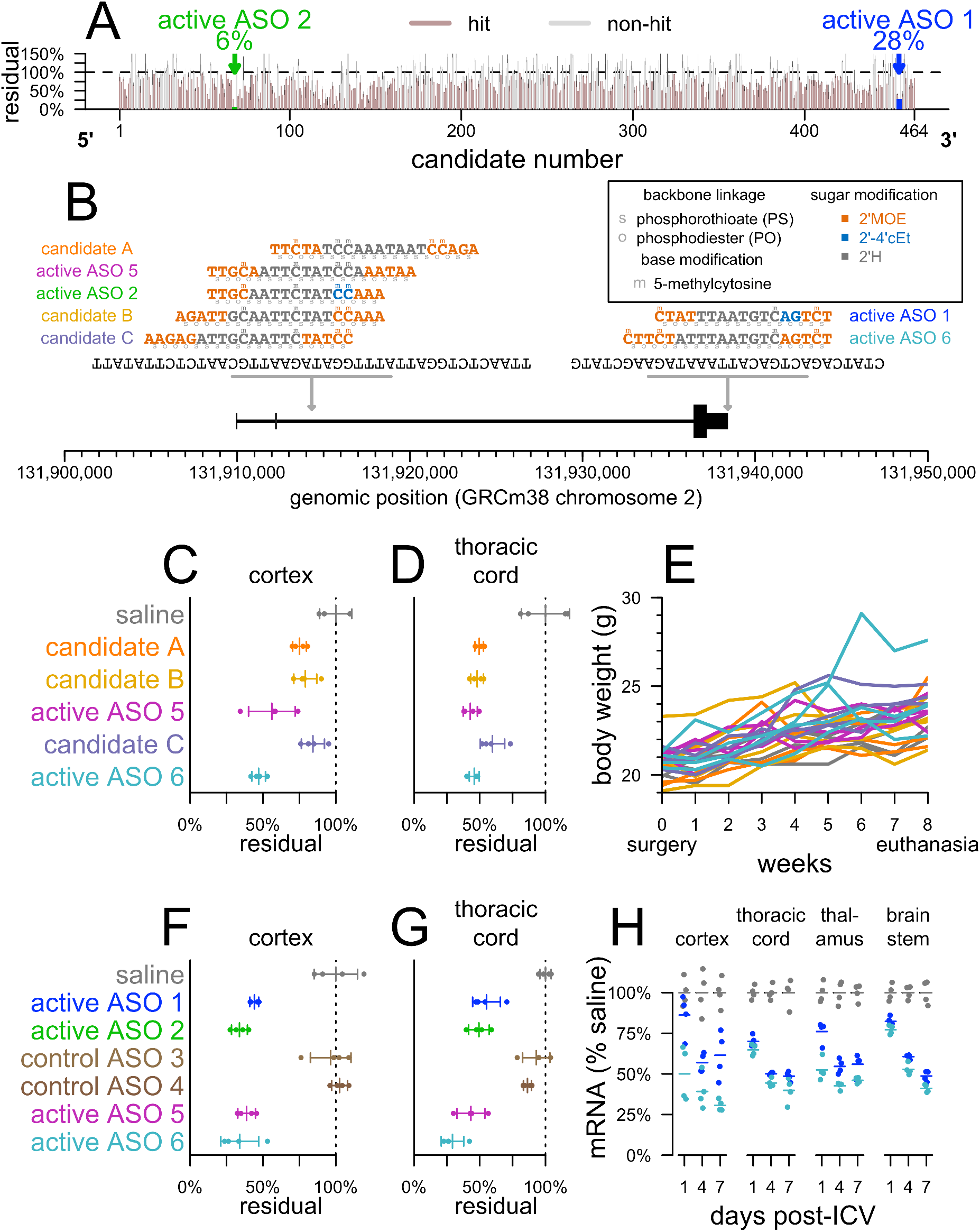
Discovery, design and characterization of ASOs used in this study. **A)** 464 ASO candidates spanning the Prnp RNA sequence were screened in HEPA 1-6 cells, and Prnp RNA was quantified as previously described [28]. 305/464 (66%) of candidates screened in cells were “hits” with a 95% confidence interval upper bound (based on N=2 replicates) of <100% of untransfected controls. **B)** The position, sequences and chemistries of previously reported active ASOs (1 and 2) [28], modified ASOs designed for the present study (5 and 6), and runner-up ASO candidates from design efforts undertaken for the present study (A, B and C). **C and D)** Groups of N=4 animals received a single 700 μg dose of the indicated treatment and ipsilateral cortex (**C**) or thoracic cord (**D**) mRNA was analyzed by qPCR 8 weeks later. **E)** Body weight trajectories for animals shown in panels C and D, over the 8 weeks between dosing and tissue analysis. **F-G)** Groups of N=4 animals received a single 500 μg dose of the indicated treatment and ipsilateral cortex (**F**) or thoracic cord (**G**) mRNA was analyzed by qPCR 1 week later. **H**) Groups of N=4 animals received a 500 μg dose of the indicated ASO and ipsilateral cortex, thoracic cord, ipsilateral thalamus, or brainstem were analyzed by qPCR 1, 4, or 7 days later.

**Table 1.** Compounds used in this study. ASOs 1, 2, 3, 4, and 6 have been previously described (58, 86). Color code for ASO chemical modifications: black = unmodified deoxyribose (2′H). orange = 2′ methoxyethyl (MOE). blue = 2′-4′ constrained ethyl (cET). Unmarked backbone linkages = phosphorothioate (PS); linkages marked with o = normal phosphodiester (PO). mC = 5-methylcytosine.

| ASO | sequence and chemistry | target |
| --- | --- | --- |
| active ASO 1 | mCToAoTTTAATGTmCAoGoTmCT | *Prnp* 3'UTR |
| active ASO 2 | TTToGomCAATTmCTATmComCoAAA | *Prnp* intron 2 |
| control ASO 3 | mComGomCTATAmCTAATomCoATAT | none |
| control ASO 4 | mCmCoToAoTAGGAmCTATmCmCAoGoGoAA | none |
| active ASO 5 | TTToGomCoAATTmCTATmCmCAoAoTAA | *Prnp* intron 2 |
| active ASO 6 | mCToTomCoTATTTAATGTmCAoGoTmCT | *Prnp* 3' UTR |

### Animals

All studies used C57BL/6N female mice purchased from Charles River Laboratories or Taconic, except for the *Prnp^+/−^* mice (45) and wild-type controls (Table 2), which were C57BL/6N of both sexes (total 38 female and 41 male), and the Tg(*Gfap*-luc) mice (46) (Figure 5), which are homozygous transgenics maintained on an FVB/N background at McLaughlin Research Institute.

**Table 2.**
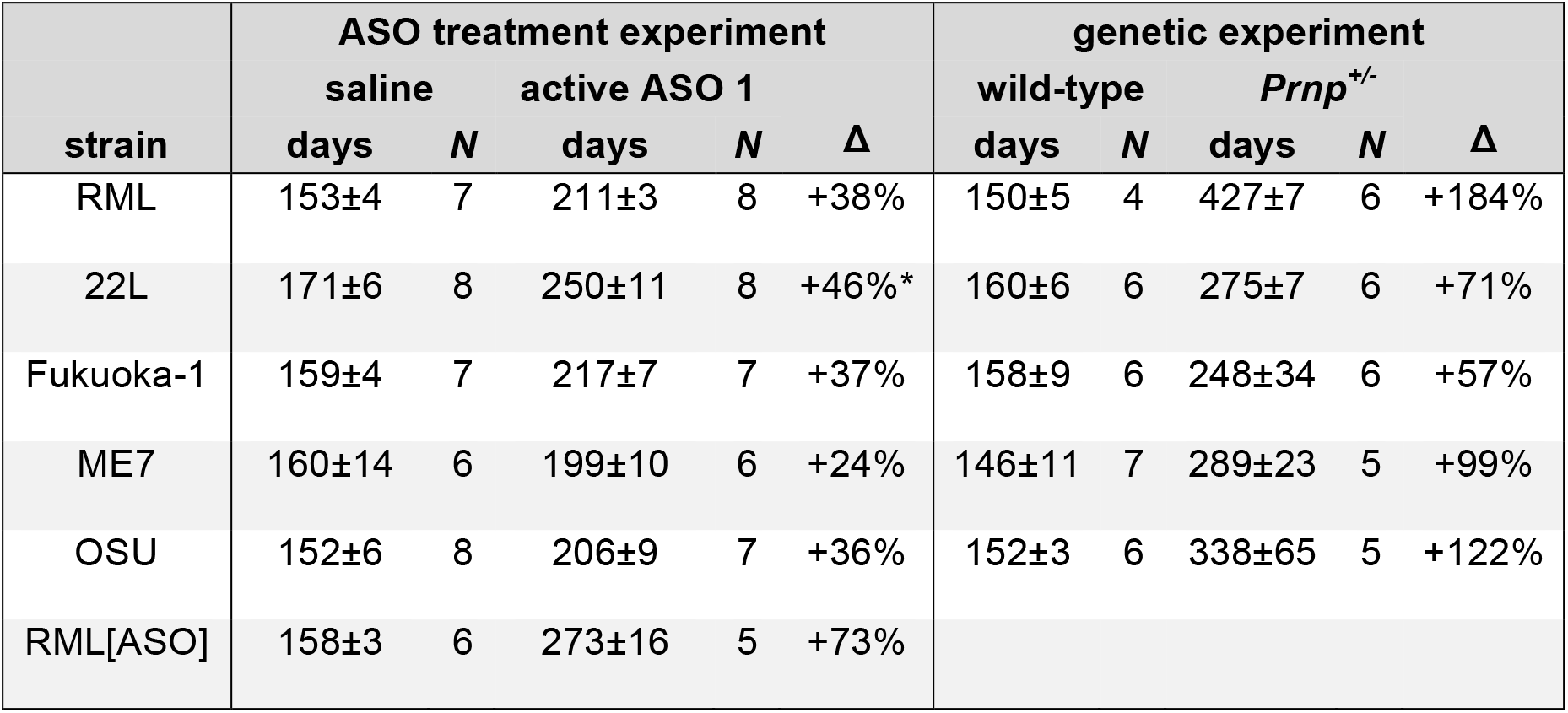
PrP lowering is effective across prion strains. Days (mean±sd) to symptomatic endpoint for animals that received two 500 μg doses of ASO vs. saline, at −14 and at 76 dpi (left, details in Figure S1), or for untreated Prnp^+/−^ vs. wild-type animals (right, details in Figure S2). For overall survival curves see Figures S1 – S2. Following established nomenclature (30), RML[ASO] denotes prions from the brains of mice infected with RML prions and treated with ASOs (see Methods). Studies conducted at the Broad Institute. *Results from repeat experiment, see Figure S1 for details.

### Prion infection

Animals were infected at age 6-10 weeks by intracerebral prion inoculation with 30 μL of a 1% brain homogenate as described (28). Briefly, brains were homogenized at 10% wt/vol in phosphate-buffered saline (Gibco 14190) in 7 mL tubes with zirconium oxide beads (Precellys no. KT039611307.7) using three 40 sec high pulses in a tissue homogenizer (Bertin EQ06404-200-RD000.0), diluted to 1% wt/vol, irradiated on dry ice at 7.0 kGy, extruded through progressively smaller-gauge blunt needles (Sai infusion B18, B21, B24, B27, B30), transferred to 2mL amber sealed sterile glass vials (Med Lab Supply), and then loaded into disposable syringes with 31G 6mm needles (BD SafetyGlide 328449). Animals were anesthetized with 3.0-3.5% isoflurane, received prophylactic meloxicam for analgesia and application of povidone/iodine as a disinfectant, and were freehand inoculated between the right ear and midline. The needle was withdrawn after three seconds and animals recovered from anesthesia in their home cages. Prion-infected brains for inoculation were supplied by co-investigators GAC (RML), KDH (Fukuoka-1 and 22L), HW, DM, and JA (ME7), and JYM (OSU). RML[ASO] brain homogenate was prepared from the pooled brains of three RML-infected animals that had received two 500 μg doses of active ASO 1 and succumbed to prion disease at 264, 270, and 270 dpi (28).

### ASO administration

ASOs were administered into CSF by bolus stereotactic ICV injection as described (28). Briefly, animals were anesthetized with 3.0-3.5% isoflurane, heads were shaved and swabbed with povidone/iodone, and prophylactic meloxicam analgesia was administered. Animals were placed in stereotaxis (ASI Instruments, SAS-4100), with 18° ear bars in ear canals and incisors in the mouse adapter tooth bar, adjusted to −8mm to level the bregma and lambda landmarks. Periosteum was scrubbed with sterile cotton-tipped applicators to reveal bregma following a ~1 cm scalp incision. Hamilton syringes (VWR 60376-172) fitted with 22-gauge Huber needles (VWR 82010-236) were filled with 10 μL of sterile saline (Gibco 14190) with or without ASO (diluted from 100 mg/mL). The needle was aligned to bregma and then moved 0.3 mm anterior, 1.0 mm right. The needle was then advanced ventral (downward) either 3.0 mm past where the bevel disappeared into the skull or 3.5 mm past where the tip of the needle first touched the skull. The liquid was ejected over ~10 seconds and the needle withdrawn 3 minutes later under downward pressure on the skull with a cotton-tipped applicator. Incisions were sutured (Ethicon 661H) with a horizontal mattress stitch. Animals recovered from the anesthesia in their home cages on a warming pad.

### qPCR

qPCR was performed as described (28) using primers *Prnp* forward: TCAGTCATCATGGCGAACCTT, reverse: AGGCCGACATCAGTCCACAT, and probe: CTACTGGCTGCTGGCCCTCTTTGTGACX; *Ppia* forward: TCGCCGCTTGCTGCA, reverse: ATCGGCCGTGATGTCGA, and probe: CCATGGTCAACCCCACCGTGTTCX. *Prnp* RNA levels were normalized to *Ppia* as a housekeeping gene and then to the mean of saline-treated controls.

### Neurofilament light quantification

Submandibular bleeds were collected with a 5mm sterile lancet (Braintree Scientific GR5MM) into a microtainer heparin blood tube (BD 365965). Tubes were inverted several times, placed on ice, and then spun at 6000 rpm for 12 min at 4°C. Plasma was transferred to a fresh cryotube and stored at −80°C until analysis. Plasma was diluted 1:4 with sample diluent and NfL was quantified using the Ella microfluidic ELISA platform (ProteinSimple) according to manufacturer’s instructions.

### Bioluminescence imaging

Each Tg(*Gfap*-luc) animal was given 5 mg (100 μL of 50 mg/mL) D-luciferin (GoldBio) in saline by intraperitoneal injection. After ~7 minutes to permit luciferin biodistribution plus ~7 minutes for 3.5% isoflurane induction, each animal was positioned into a Lumina II in vitro imaging system (IVIS; Perkin Elmer) with nosecone isoflurane maintenance and imaged for 1 minute before returning to its cage. At each session, three control Tg(*Gfap*-luc) animals were imaged to test luciferin and equipment: two mice that received intraperitoneal lipopolysaccharide (LPS; positive control causing brain gliosis), and one mouse that received saline (negative control) 16h prior. Data for a single region of interest (ROI), defined based on an LPS positive control animal, were extracted using Living Image Software 4.5 (Perkin Elmer). Bioluminesence was measured in photons per second emitted from one square centimeter of tissue radiating into a solid angle of one steradian (sr) — photons/sec/cm^2^/sr, also called radiance units or simply photons. This calibrated measure controls for charge coupled device (CCD) camera settings such as F-stop, exposure, and binning, in contrast with absolute measurement of incident photons, allowing adjustment of camera settings without compromising comparability of results.

### Rotarod

Mice were seated on a rod rotating at 4 rpm in a 6-lane Rotarod apparatus (Maze Engineers). Once all mice from a single cage were properly seated, rotation was accelerated at 6 rotations per min for 5 min, and then held constant at 34 rpms for another 5 min. Latency to drop was recorded, in seconds, with a maximum score of 600 seconds if the mouse did not fall or ease itself off the rod. At each time point, the mice underwent 9 trials (3 trials per day over 3 days), with trials 1-3 considered to be spent learning the task and trials 4-9 included in analysis.

### Disease monitoring and endpoints

At the Broad Institute, animals were checked for general health daily and subjected to detailed monitoring once weekly beginning at 90 dpi and thrice weekly beginning at 120 dpi. In these monitoring sessions, animals were weighed, and scored 0 or 1 for each of eight behavioral tests: scruff / poor grooming, poor body condition, reduced activity, hunched posture, irregular gait / hindlimb weakness, tremor, blank stare, and difficulty righting. Detailed observational criteria and performance statistics for these tests are provided in Table S1. Nest-building was rated for both cotton square nestlets (Ancare) and Enviro-dri® packed paper (Shepherd) on a scale of 0 = unused; 1 = used/pulled apart, but flat; 2 = pulled into a three-dimensional structure. Cotton and paper scores were averaged to yield a combined score. Animals were group housed. The rare instances of cages shared by animals of different treatment cohorts were excluded from nest analyses. Animals were euthanized by CO_2_ inhalation when they met pre-defined endpoint criteria. Terminal endpoint criteria, intended to catch mice just shortly before disease progressed naturally to death, were defined initially as body condition score <2, body weight loss ≥20% from baseline, inability to reach food or water, severe respiratory distress, or severe neurological deficits (Figure 2D-F), and later refined to simply body weight loss ≥15% from baseline or inability to reach food or water (Figure 7). Symptomatic endpoint criteria, intended to catch mice at an advanced disease stage but before terminal illness, were defined as ≥5 of the 8 pre-defined symptoms being observed at two consecutive monitoring sessions, or body weight loss ≥15% from baseline, body condition score ≤2, or inability to reach food or water (Figures 3 and 6 and Tables 2 and 3). At the McLaughlin Research Institute, mice were monitored for diverse neurological and non-neurological health indicators and SHIRPA phenotypes (47) (Table S3) in the natural history study (Figure 4), and checked for general health and weight in other studies (Figure 5); they were euthanized at ≥20% body weight loss from baseline, inability to reach food or water, or moribund status.

**Figure 2.**
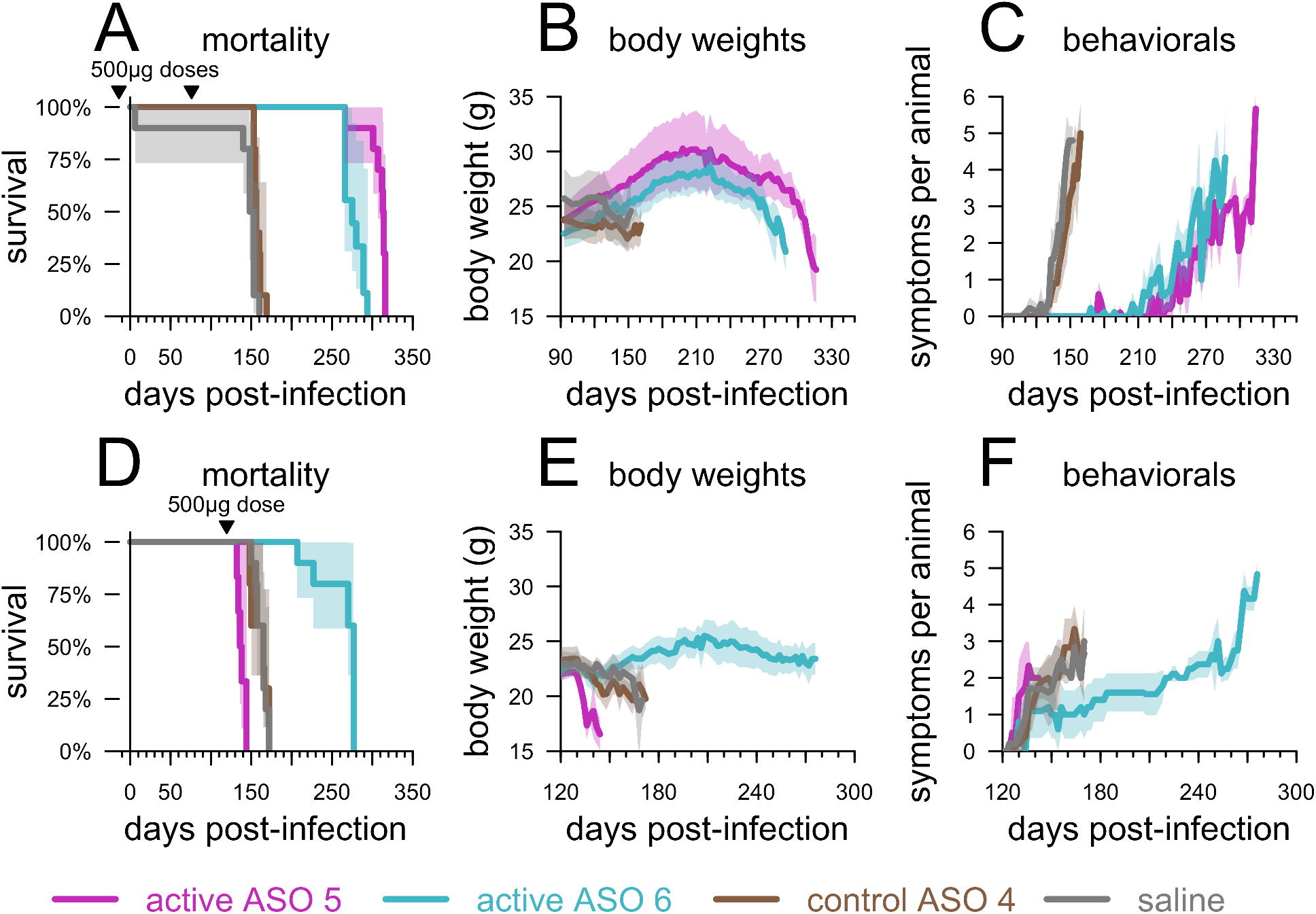
Replication of early and late treatment efficacy of ASOs. Survival **(A,D)**, body weights **(B,E)**, and symptom trajectories **(C,F)** of mice treated with ASOs prophylactically (−14 and 76 dpi) **(A-C)** or at 120 dpi **(D-F)**. Shaded areas represent 95% confidence intervals.

**Figure 3:**
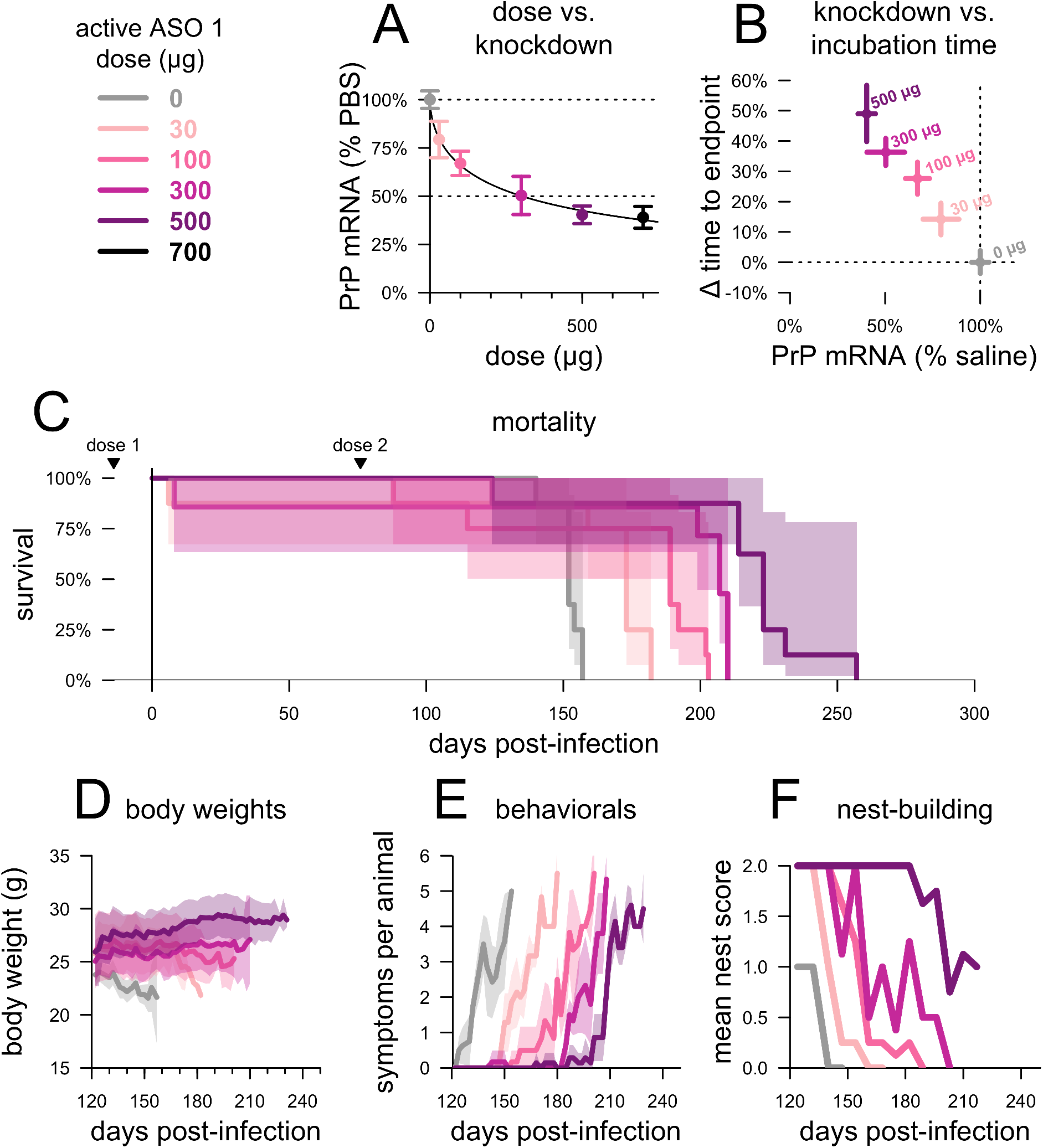
Relationship between degree of PrP lowering and therapeutic benefit. **A)** Dose versus ipsilateral cortical PrP mRNA knockdown determined by qPCR at 2 weeks post-treatment and normalized to the mean of saline-treated, non-infected animals, N=3 per group, **B)** PrP mRNA knockdown (from panel A) versus time to symptomatic endpoint in groups of N=8 prion-infected animals receiving two injections of the indicated dose, at −14 and 76 dpi, and, for the same animals, **C)** overall mortality, **D)** body weights normalized to each mouse’s individual weight at 122 dpi, **E)** mean symptom count per animal, and **F)** mean nest score. Studies conducted at the Broad Institute. Shaded areas represent 95% confidence intervals.

**Figure 4.**
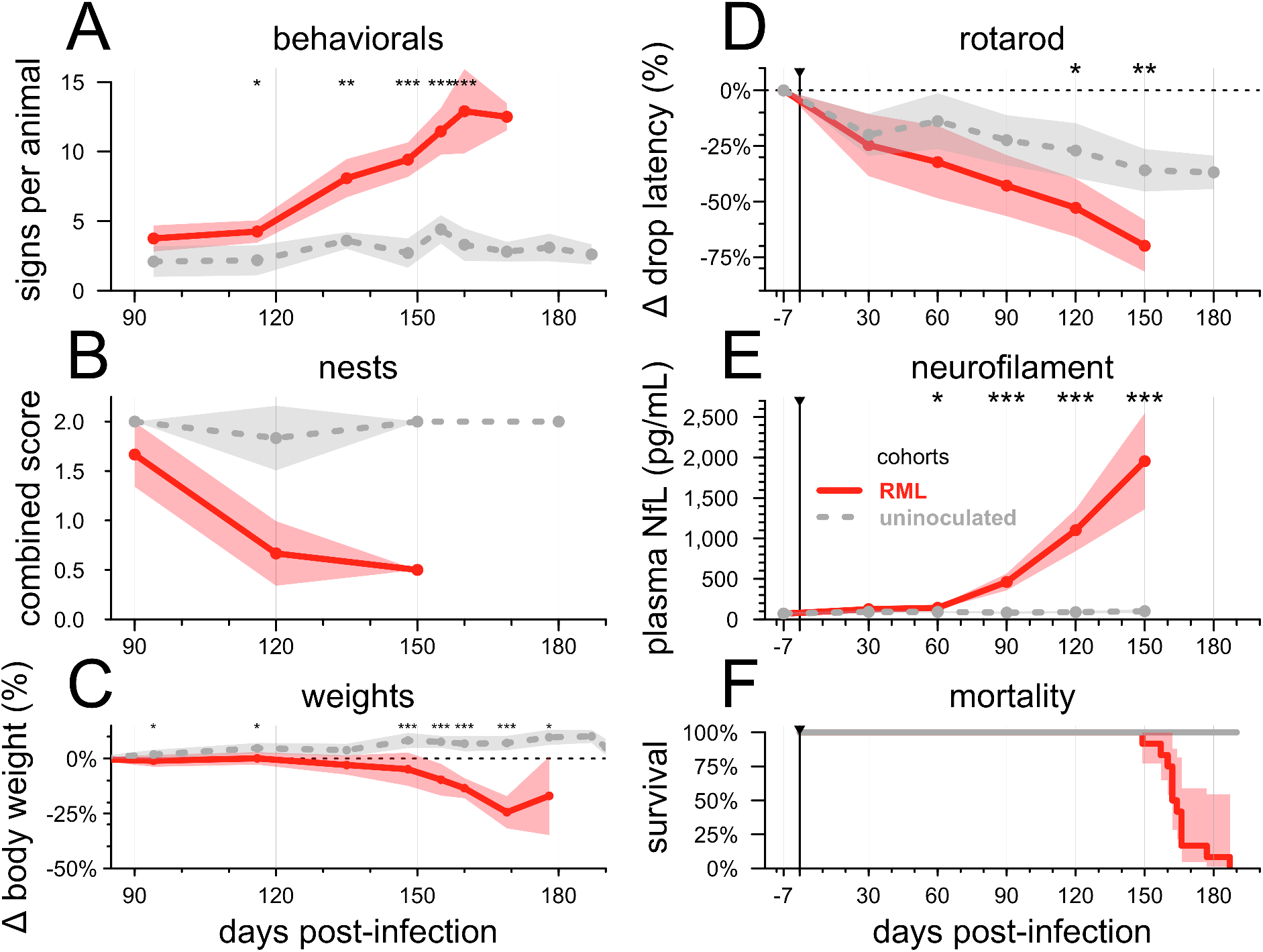
Natural history of RML prion infection. N=12 mice infected with 30 μL of a 1% RML brain homogenate versus N=12 uninoculated controls. In panels A-E, lines represent means, shaded areas represent 95% confidence intervals of the mean, and dots represent assessment timepoints. Nominal statistical significance thresholds (two-sided Kolmogorov-Smirnov test) are displayed as: * P < 0.05, ** P < 0.01, *** P < 0.001. **A)** symptom accumulation (see Figure S4 and Table S3 for details), **B)** nest-building scores, **C)** weight change relative to each animal’s 78 dpi baseline (see raw individual weights in Figure S5A)†, **D)** rotarod performance relative to each animal’s −7 dpi baseline (see raw individual latencies in Figure S5B), **E)**, plasma NfL (see raw individual NfL trajectories in Figure S5C), and **F)** overall mortality. †In panel C, prion-infected animals that reached endpoint between planned assessments and were weighed a final time prior to euthanasia are grouped together with animals at the next planned assessment timepoint — for example, animals that reached endpoint at 166 dpi are averaged into the 169 dpi timepoint. Studies conducted at McLaughlin Research Institute.

**Figure 5.**
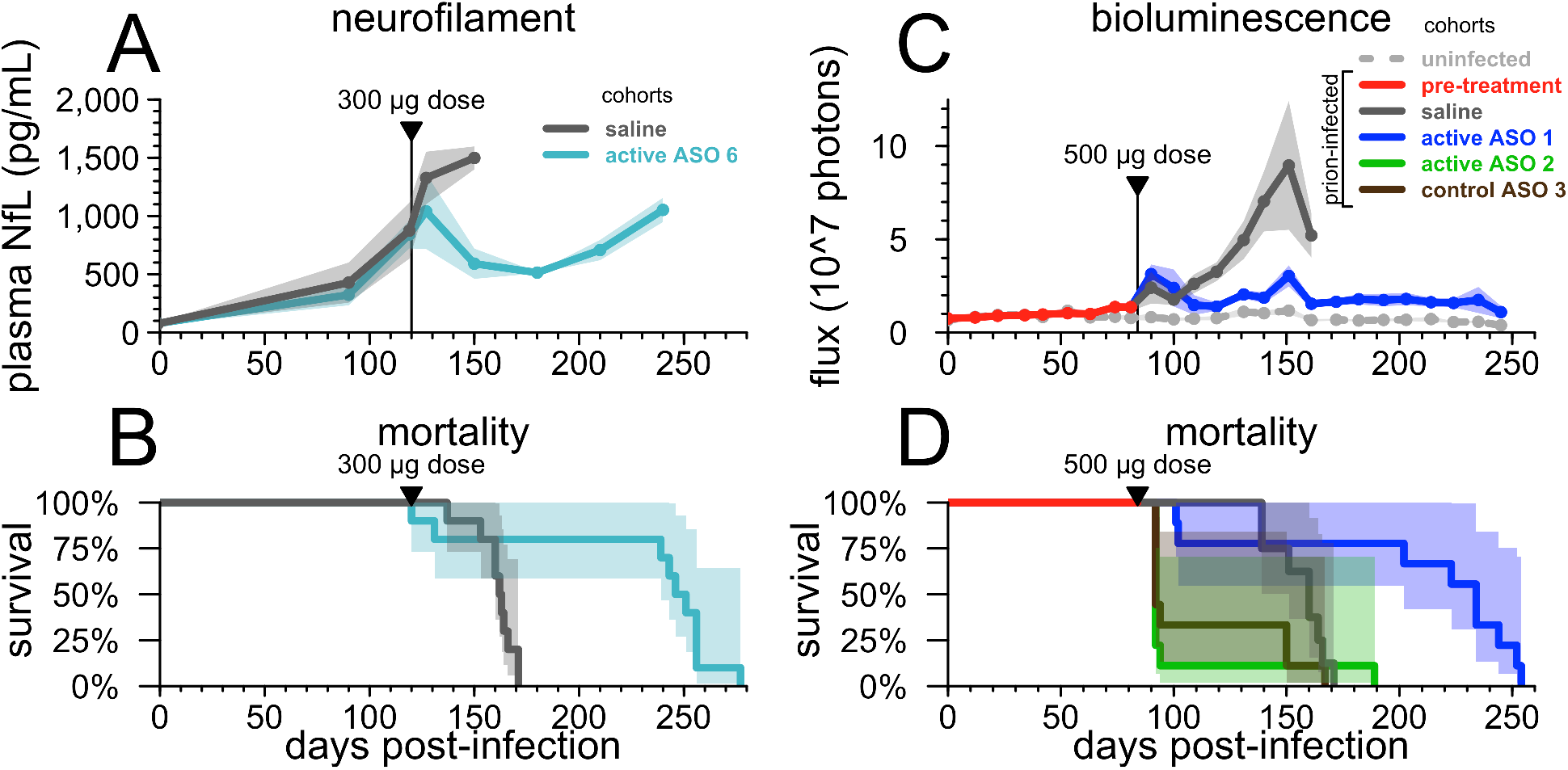
Response of neuronal damage and astrocytosis biomarkers to ASO treatment at a pathological timepoint. **A)** plasma NfL and **B)** survival in wild-type mice infected with prions and dosed at 120 dpi, a timepoint at which the natural history study (Figure 4D) had indicated that NfL was dramatically elevated and rotarod performance and nest-building might be impaired. N=10 per group, of which NfL was assessed in N=10 saline-treated and N=5 active ASO 6-treated animals. **C)** live animal bioluminescence and **D)** survival in Tg(Gfap-luc) mice infected with prions and dosed at 83-84 dpi, after two consecutive imaging sessions showed elevated luminescence in the RML group compared to uninfected controls. N=9 per treatment group plus N=14 uninfected controls. Shaded areas represent 95% confidence intervals. Studies conducted at McLaughlin Research Institute.

**Figure 6.**
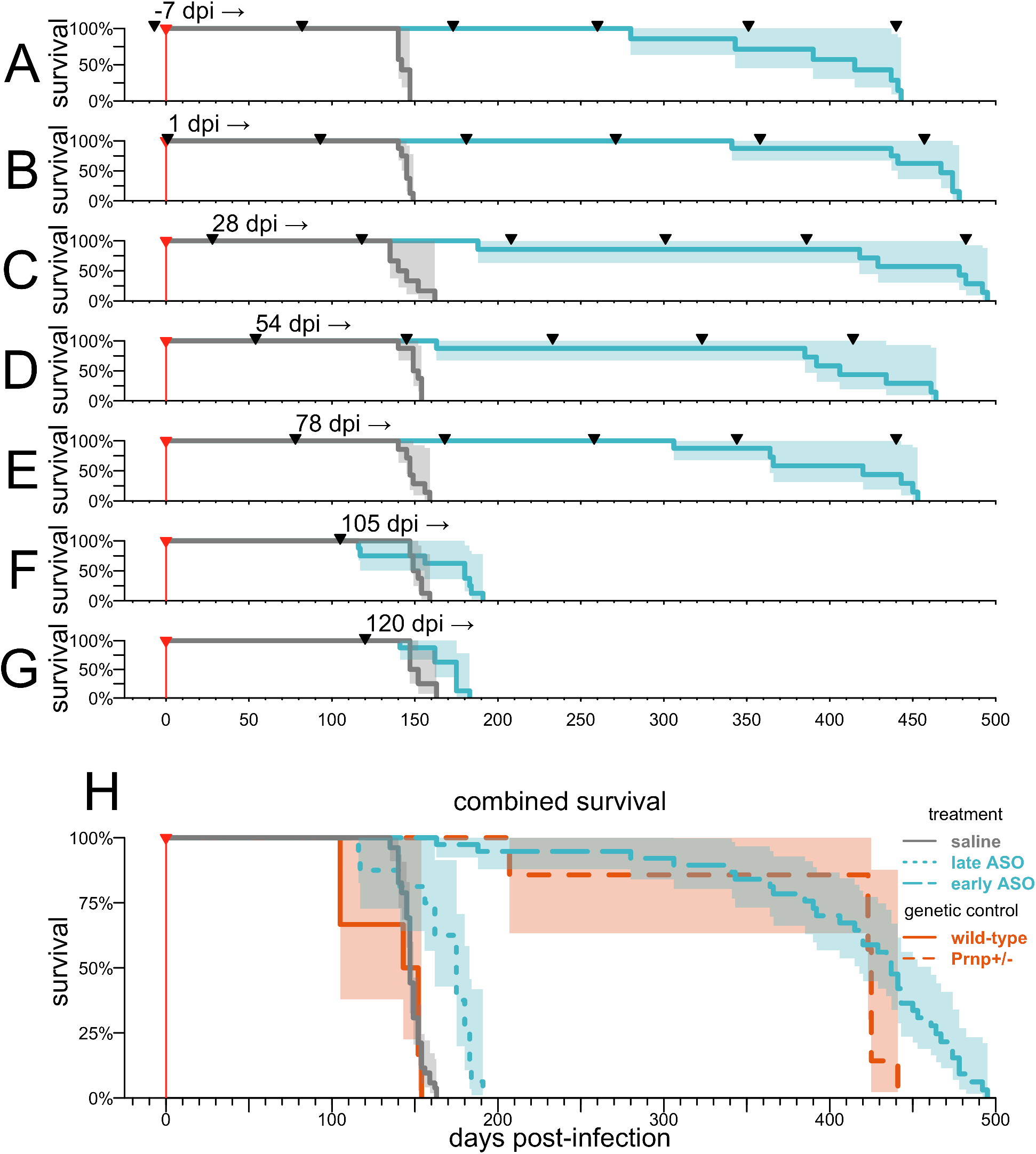
Efficacy of PrP-lowering therapy is timepoint-dependent. Groups of N=8 animals received saline or active ASO 6, chronically every ~90 days beginning at the specified timepoint. Black triangle indicated when ASO was injected. **A-G)** Survival time as a function of time of treatment initiation, **H)** combined survival curves for saline-treated mice versus mice treated with active ASO 6 at early (−7 to 78 dpi) or late (105 to 120 dpi) timepoints. Survival curves for wild-type versus Prnp+/− animals infected with RML prions shown in Table 2 and Figure S2 are reproduced here for comparison. Studies conducted at the Broad Institute. Shaded areas represent 95% confidence intervals.

**Figure 7.**
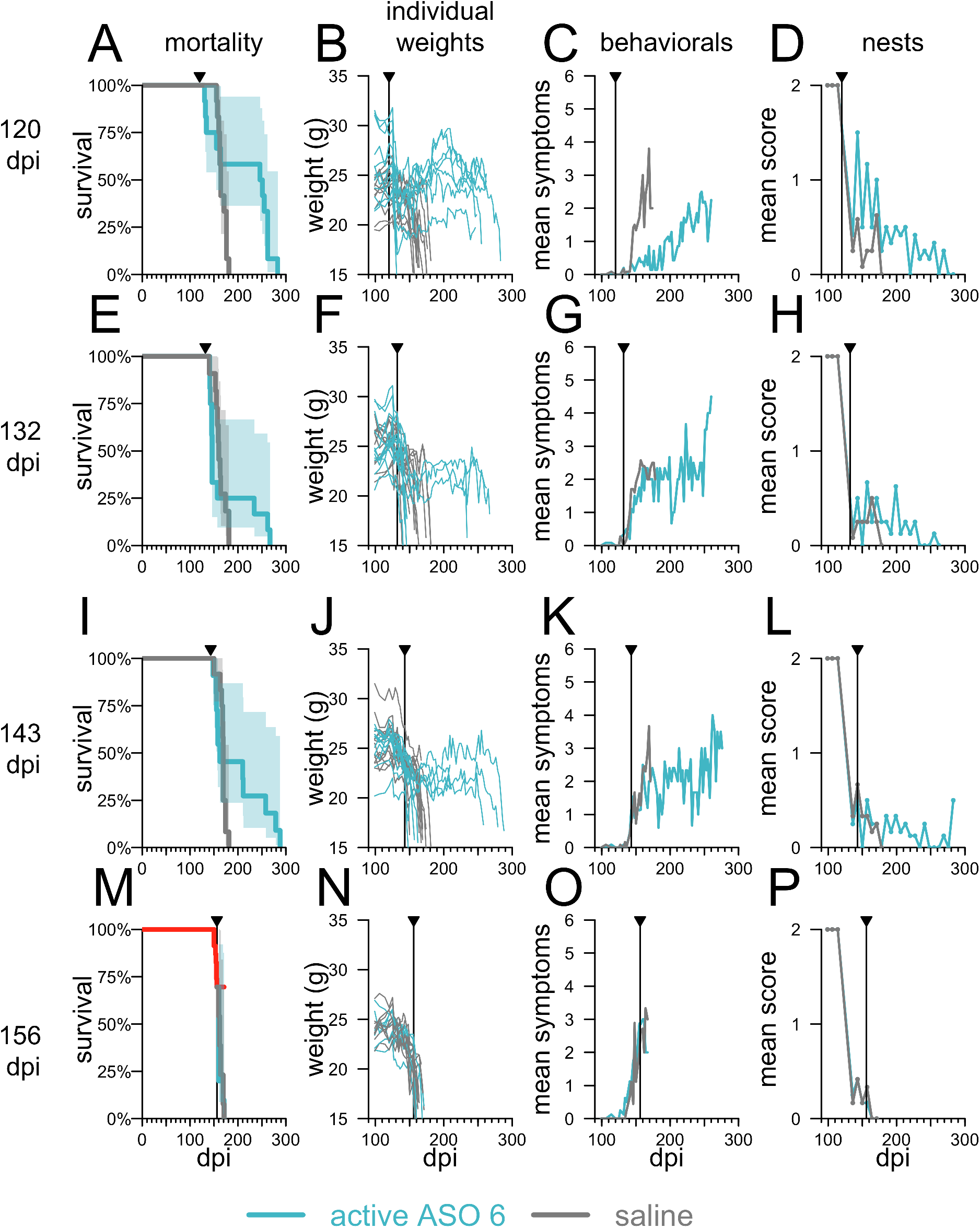
Effects of intervention at pathological and symptomatic timepoints. Animals were infected with RML prions and then received saline (N=12) or a single 500 μg dose of active ASO 6 (N=12) at the indicated timepoint. A, E, I, M) survival; B, F, J, N) individual body weight trajectories; C, G, K, O) symptom count summarized by cohort; D, H, L, P) nest-building activity summarized by cohort. For the 156 dpi timepoint, 7/23 animals (30%) reached endpoint prior to the intervention (red curve, panel M). Shaded areas represent 95% confidence intervals. Studies conducted at the Broad Institute.

**Table 3.** PrP lowering is effective in delayed intervention against multiple prion strains. Mice were infected with any of five prion strains and then treated with 500 μg ASO, or saline, at a pre-specified timepoint expected to be 80% of the way through the control group incubation period based on a previous experiment (Table 2). Actual treatment timepoints ranged from 75% - 81% of the incubation period. Overall Δ indicates mean difference in time to symptomatic endpoint counting all animals. The rightmost three columns show the number and mean survival of those ASO-treated animals that survived at least 10% longer than the mean of the saline animals for each strain. Details visualized in Figure S3. Studies conducted at the Broad Institute. *One of two cages intended for the Fukuoka-1 active cohort was lost to experimental error, resulting in a lower N for this group.

| strain | intervention<br>timepoint | | saline | | active ASO 6 | | overall<br>$\Delta$ | active ASO 6 animals<br>surviving >10% longer<br>than saline mean | | |
| --- | --- | --- | --- | --- | --- | --- | --- | --- | --- | --- |
| | dpi | relative | days | N | days | N | | proportion | days | $\Delta$ |
| RML | 123 | 75% | 164±7 | 6 | 189±54 | 8 | +15% | 3/8 | 251±17 | +53% |
| 22L | 127 | 78% | 164±5 | 8 | 235±43 | 7 | +44% | 6/7 | 251±5 | +54% |
| Fukuoka-1 | 128 | 77% | 166±14 | 8 | 255±4 | 3* | +54% | 3/3 | 255±4 | +54% |
| ME7 | 129 | 81% | 160±11 | 8 | 179±52 | 8 | +12% | 3/8 | 234±45 | +46% |
| OSU | 122 | 77% | 159±6 | 8 | 204±60 | 7 | +28% | 4/7 | 250±17 | +57% |

### Statistical analysis

Data analysis, visualization, and statistics were conducted in R 3.6.1. All statistical tests were two-sided and reported *P* values are nominal. To avoid selective reporting of only those deaths subjectively attributed to prion disease, survival curves reported herein include all causes of death except for the following: death prior to any drug treatment (meaning prior to experimental treatment group being assigned); acute deaths within 1 day post-surgery due to surgical complications; and euthanasia due to experimental error (such as incorrect dosing or inability to position animal in stereotaxis). For dose-response data (Figure 3), where incubation times within cohorts were approximately normally distributed, changes in incubation time and confidence intervals thereof were determined using t-test ratios (48) in the R `mratios` package (49). Survival outcomes in many of our other experiments were bimodal, and testing using the R `survival` package (50) revealed that these did not meet the proportional hazards assumption (51), which is required for log-rank tests. Survival differences were therefore determined by visual inspection with shaded 95% confidence intervals computed by the log transform method (52). Dose-responsiveness of target engagement was fitted using a four-parameter log-logistic regression with the R `drc` package (53). Distributions of biomarker and behavioral outcomes were compared using two-sided Kolmogorov-Smirnov tests, which do not assume normality. For the bioluminescence study (Figure 5 C-D), ASO administration was pre-specified to occur after two consecutive imaging sessions in which bioluminescence differed between infected and uninfected mice at *P* < 0.05 by a Kolmogorov-Smirnov test.

### Data availability

We have provided a public data repository at https://github.com/ericminikel/prp_lowering to enable others to reproduce our analyses. This repository contains individual-level animal data for every experimental animal, including weights, behaviorals, nest scores, biomarkers, and survival endpoints, as well as R source code to generate all of the figures and tables in this manuscript from those raw data.

### Study approval

All experiments were conducted under approved Institutional Animal Care and Use Committee protocols (Broad IACUC 0162-05-17, Ionis IACUC P-0273, and McLaughlin IACUC 2017-GAC22 / 2018-MPK29).

## Results

### Therapeutic benefit and mechanism of action replicate across ASO chemistries

ASOs can be synthesized with diverse combinations of sugar, backbone, and other chemical modifications (54). Survival benefits in prion-infected mice have been previously demonstrated for three PrP-targeting sequences with two chemical formulations (22, 28). The first reported PrP-lowering ASO had 2’-*O*-methoxyethyl (MOE) wing modifications and a straight phosphorothioate (PS) backbone (22). Subsequently, we identified through cellular screening two potently PrP-lowering ASOs with mixed MOE and 2’-4’ constrained ethyl (cEt) wings, and a mixed backbone containing PS as well as normal phosphodiester (PO) linkages (ASOs 1 – 2, Table 1, Figure 1A) (28). Previously reported *in vivo* studies with ASOs 1 and 2 and a chemically matched, non-targeting control confirmed target RNA and protein reduction *in vivo*, and showed that lowering of the RNA was required for beneficial effect in prion-infected mice, suggesting that the oligonucleotides were acting through an RNase-H mediated mechanism (28).

Motivated by the desire to additionally test the ASO chemistry now in clinical trials for Huntington’s disease and Amyotrophic lateral sclerosis (ALS) with SOD1 mutations (25, 55), we undertook optimization at the binding sites of active ASOs 1 and 2 to design and synthesize a set of ASOs with mixed PS/PO backbones and a 10-base deoxynucleotide gap flanked on each end with 2’O-methoxyethyl (MOE) modified nucleotides (Table 1, Figure 1B). As before, newly designed ASOs were prioritized based on potency of prion protein RNA knockdown (Figure 1C-D) *in vivo*. Over eight weeks of post-dose monitoring, there were no findings in weekly neurological exams,,and behavioral observations and body weight gain trajectories were comparable to those of saline-treated control animals (Figure 1E). Selected compounds (ASOs 5 and 6) achieved similar levels of target engagement as those previously reported, with active sequences reducing cortical PrP RNA by approximately half within one week after a 500 μg dose while both the previously reported and a new, chemistry-matched, control ASO were confirmed inactive (Figure 1F-G). Compared to the original active ASOs, newly designed compounds showed comparable time to effect (Figure 1H). Following this initial characterization, we sought to replicate previous results by evaluating the new active and inactive ASOs in a prion disease model.

We studied the efficacy of ASOs in intracerebrally prion-inoculated mice in experiments variously utilizing either a symptomatic endpoint (euthanasia upon observation of five pre-specified neurological symptoms; see Methods) or a more advanced terminal disease endpoint (euthanasia upon 15-20% body weight loss or inability to reach food and water; see Methods). These paradigms respectively allow for early halting of experiments when animals have become moderately ill, or for the potential to observe changes in the rate of symptomatic progression towards end-stage disease.

In a prophylactic experiment as previously described (28), intracerebroventricular (ICV) ASO treatments were administered at 14 days prior to and again at 76 days post-infection (dpi) with Rocky Mountain Lab (RML) prions (56), a widely used laboratory prion strain (57). Groups of N=10 C57BL/6N mice received two 500 μg doses of active ASO 5, active ASO 6, control ASO 4, or saline by stereotactic ICV injection. Active ASOs 5 and 6 closely replicated the survival benefit reported with active ASOs 1 and 2 (28), delaying symptomatic endpoint by 108% and 80% respectively compared to saline (median 314 and 270 vs. 150 dpi) (Figure 2A, Table S2). These PrP-targeting ASOs delayed onset of disease as reflected in weight loss (Figure 2B) and symptom accumulation (Figure 2C) in treated animals. In a delayed treatment experiment mirroring that reported previously (28), a single 500 μg bolus dose was administered at 120 dpi, or ~72% of the time to terminal disease endpoint. This terminal endpoint was delayed by 68% for active ASO 6 (median 277 vs. 165 dpi; Figure 2D, Table S2), with all mice surviving beyond the point when all of the saline-treated animals had died, while weight loss was partially reversed and symptom accumulation attenuated (Figure 2E-F). Active ASO 5 was not tolerated at this timepoint (Figure 2D, Table S2), replicating the ASO-specific, disease stage-dependent toxicity reported previously (22, 28). Across both prophylactic and delayed treatment paradigms, non-targeting control ASO 4 conferred no survival benefit (Figure 2A, 2D), replicating control ASO 3 results (28) and confirming PrP lowering as the mechanism of action by which ASOs antagonize prion disease (28, 58). In both of these experiments, with blinded assessments (see Methods), we recapitulated our previous findings, demonstrating that ASO-mediated PrP lowering extended survival and delayed disease course, in both prophylactic and delayed treatment paradigms. Given the comparable results across tool compounds of different chemistries, active ASOs 1 and 6 were used interchangeably in the experiments that follow.

### Dose-responsive benefits to PrP-lowering

We next investigated the minimum level of PrP suppression sufficient to confer benefit in prion-inoculated mice. Toward this end, we injected ASO1 into wild-type, uninfected mice at 6 doses (0-700 μg), and found a dose-dependent reduction in *Prnp* mRNA in the cortex at 2 weeks post-dose was dose-dependently lowered, with residual RNA levels ranging from 79% at the 30 μg dose to 39% at the 700 μg dose compared to vehicle-treated animals (Figure 3A). As target engagement at the 500 and 700 μg doses was not significantly different, the 0 through 500 μg doses were selected for a survival study in RML prion-infected mice per prophylactic paradigm described above (two doses, at −14 and 76 dpi) utilizing a symptomatic endpoint assessed by blinded raters. Across doses of 0 (saline), 30, 100, 300, or 500 μg of active ASO 1, *Prnp* RNA reduction tracked with incubation time in animals that ultimately succumbed to prion disease (Figure 3B), with a significant increase in time to symptomatic endpoint even at 21% knockdown (median 173 vs. 152 dpi at 30 μg, *P* = 0.002, two-sided log-rank test). Across all doses, overall survival was increased in step with knockdown (Figure 3C, Table S2) and attendant delays in weight loss (Figure 3D), accumulation of prion disease symptoms (Figure 3E), and decline in nest-building (Figure 3F) suggested that at all doses tested, the treatment had extended healthy life. Thus, dose-dependent PrP lowering translated to dose-dependent benefit in prion disease, with as little as 21% RNA knockdown extending survival.

### Efficacy of PrP lowering across prion strains

As all prion strains share the common substrate of PrP, we hypothesized that reduction of PrP, by either genetic or pharmacologic means, would effectively modify prion disease across strains. To test this hypothesis, we challenged mice with five different previously characterized mouse-adapted laboratory prion strains of diverse origins: RML (adapted from goat scrapie) (56), 22L (sheep scrapie) (59), ME7 (sheep scrapie) (60), Fukuoka-1 (human P102L GSS) (61), and OSU (synthetic) (62).

In the pharmacological treatment arm, groups of mice infected with these prion strains received 500 μg ASO 1 at −14 and 76 dpi or saline (N=8 per treatment per strain). In the genetic control arm, heterozygous ZH3 PrP knockout (45) *(Prnp^+/−^*) or wild-type mice (N=8 per genotype per strain) were inoculated with the same five prion strains listed above without pharmacologic intervention. Mice in both arms were followed to a symptomatic endpoint by blinded raters. Across strains, disease was delayed and survival extended in animals with reduced PrP, whether the reduction was ASO-mediated (Table 2, Figure S1) or genetic (Table 2, Figure S2). Survival time response to ASO treatment across strains ranged from +24% to +46%, while the increase in survival time due to heterozygous PrP knockout ranged from +57% to +184% (Table 2), with differences among strains reflected in overall mortality and in trajectories of body weight loss, symptom accumulation and nest-building (Figure S2). Overall, prophylactic PrP lowering by genetic or pharmacologic means proved effective against all five strains tested.

To test whether ASO treatment gives rise to drug-resistant prion strains, we prepared brain homogenate from terminally sick, RML prion-infected, active ASO 1-treated animals included in a previous experiment (28) (see Methods). Groups of N=8 mice inoculated with this prion isolate, termed RML[ASO] following established nomenclature (30), received two doses of 500 μg active ASO 1 or saline per the described prophylactic paradigm. Active ASO 1 retained its efficacy in this paradigm, delaying symptomatic endpoint by 74% (Table 2, Figure S1), similar to the 61% delay in the experiment from which the RML[ASO] isolate was sourced (28), suggesting that ASO treatment does not give rise to drug-resistant prion strains.

We next sought to compare the effect of PrP-lowering treatment across multiple strains in delayed treatment. We chose intervention timepoints for each strain estimated to be after ~80% of the incubation time had elapsed, based upon the previous experiment (Table 2), thus roughly corresponding to the 120 dpi timepoint where we and others observed efficacy against RML prions (Figure 2 and ref. (28)). At the chosen timepoint (122-129 dpi), each group of N=8 mice received one dose of 500 μg of active ASO 6 or saline and was followed to a symptomatic endpoint, again by blinded raters (see Methods). At this timepoint, active ASO 6 remained effective against all five prion strains (Table 3). In terms of increase in mean survival time, the ASO appeared highly effective against some strains and marginally effective against others (Table 3), however, inspection of survival curves (Figure S3A) revealed that differences were driven not by differences in maximum survival time, but by the proportion of ASO-treated animals that outlived their saline-treated counterparts. Accordingly, for each strain, we applied a cutoff of survival 10% beyond the mean of saline-treated controls (Table 3, right panel), corresponding to 1.96 standard deviations of control survival, when 95% of control animals would be expected to have reached endpoint. The differences in the proportions of ASO-treated animals crossing this threshold were not significantly different between strains (*P* = 0.80, two-sided Fisher exact test) and, among these animals, the overall mean survival time increase was similar across strains (+46% to +57%). Across strains, for treated animals that outlived controls, body weights declined initially and then partly rebounded (Figure S3B), first symptoms emerged on a timeline similar to controls but further symptoms accumulated more slowly (Figure S3C), and nest building was somewhat impaired in the treated animals, with some variability between strains (Figure S3D). Overall, efficacy of late PrP-lowering treatment was confirmed across all five strains tested.

### Natural history of RML prion infection

In order to establish the pathological context of different treatment timepoints, we endeavored to systematically map biomarker, weight, and behavioral changes onto the incubation period by comparing N=12 RML prion-infected mice and N=12 uninoculated controls. Rotarod performance, an early sign in some prion models (63), and neurofilament light (NfL) in blood, an early sentinel biomarker of more slowly progressive neurodegenerative diseases in both mice (64) and humans (65, 66), were evaluated at −7 dpi and every 30 days following inoculation. Weights, nest-building activity and a battery of symptomatic and behavioral observations (Table S3) were evaluated as the animals approached terminal endpoint.

Overall, group-wise symptomatic changes became apparent at approximately 120 dpi (Figure 4). Across 40 symptomatic and behavioral observations conducted (Table S3), the mean number of observations with score >0 became nominally elevated in RML mice at 116 dpi and unambiguously elevated by 135 dpi (*P* = 0.017 and *P* = 0.0010 respectively, two-sided Kolmogorov-Smirnov tests, Figure 4A). No individual observation measure showed any earlier sensitivity, with clear changes only at 135 dpi in abnormal activity level (slow), no balance on bar, and tail suspension: poor or no splay (Figure S4). Nest-building was impaired in all prion-infected cages by 120 dpi, though with just N=3 cages per group the significance of this remained ambiguous (*P* = 0.10, two-sided Kolmogorov-Smirnov test, Figure 4B). Weight loss, relative to each animal’s baseline weight, achieved nominal significance in some but not all weighing sessions from 94 dpi onward, but became unambiguous only at 148 dpi (Figure 4C).

Rotarod performance in prion-infected mice, normalized to each mouse’s own baseline, began to show nominal decline at 120 dpi (*P* = 0.028, Figure 4D) strengthening by 150 dpi (*P* = 0.0024, Figure 4D). Even as these differences became apparent on a group-wise basis, distributions of both weights and rotarod latencies overlapped until some animals began to reach endpoint (Figure S5A-B). In contrast to these symptomatic measures, molecular evidence of pathology was detectable far sooner. Plasma NfL was nominally increased in prion-infected mice at 60 dpi (*P* = 0.015, two-sided Kolmogorov-Smirnov test), with a subset of mice elevated while the distributions still overlapped (Figure S5C). By 90 dpi, plasma NfL levels showed clear elevation in prion-infected mice, with non-overlapping distributions (Figure 4E and S5C) preceding frank symptoms. All changes grew in magnitude until the prion-infected mice reached endpoint at a median of 163 dpi (Figure 4F).

### Biomarker response in mice treated at a pathological timepoint

Having characterized the time course of pathology, we evaluated whether and how biomarkers of pathology respond to PrP-lowering treatment. To evaluate NfL response to treatment, groups of N = 10 mice were inoculated with RML prions, and received a single ICV bolus dose of ASO 6 or saline at 120 dpi. Plasma NfL was quantified from bleeds taken at −1 dpi, 90 dpi, 119 dpi (one day pre-dose), 127 dpi (one week post-dose), and then every 30 days from 150 dpi onward. As expected, plasma NfL levels steadily rose through terminal illness in saline-treated animals (Figure 5A). In contrast, by 30 days after ASO treatment, plasma NfL levels fell significantly in ASO-treated mice compared to the immediate pre-dose timepoint, suggesting a reversal of pathology driving the 53% increase in survival time (median 248 vs. 162 days, Figure 5B, Table S2). NfL began to rebound ~90 days post-treatment, coincident with expected waning of the pharmacodynamic effect of ASOs (28) (Figure 5A). This experiment provided biomarker evidence that ASO-mediated PrP lowering can reverse pathology after disease-associated changes have begun to occur. To our knowledge, this is the first time pharmacological reversal of a translatable biomarker of disease has been demonstrated in a prion-infected animal.

Reactive gliosis associated with increased expression of the astroglial intermediate filament gene *Gfap* has been previously established as one of the earliest neuropathological changes in prion-infected mice (67). Using Tg(*Gfap*-luc) mice (46), which express luciferase under the *Gfap* promoter, it is possible to track the progression of gliosis by live animal bioluminescence imaging (BLI) throughout the course of prion disease (68) and to obtain time-series data on the effect of drug treatment(31). To evaluate astroglial proliferation, we imaged N=36 Tg(*Gfap*-luc) RML prion-infected and N=14 uninfected mice by BLI every 7-11 days, and pre-specified that a single 500 μg dose of ASO 1, 2 or 3 would be administered after two consecutive imaging sessions showed a nominally significant (*P* < 0.05 by a two-sided Kolmogorov-Smirnov test) difference in BLI between infected and uninfected mice. Significant differences were observed at 73 and 81 dpi, triggering the ASO injections to be performed at 83-84 dpi (Figure 5C).

Consistent with our previous report (28), ASO 2 and control ASO 3 were poorly tolerated at a pathological timepoint: 8/9 animals treated with active ASO 2 and 6/9 treated with control ASO 3 died or were euthanized 8-11 days post-surgery. In the active ASO 1 cohort, 2/9 animals also died 17-19 days post-surgery. Across treatment groups, all mice that survived the three-week period after surgery eventually developed progressive neurological signs consistent with prion disease, although half (9/18) of these mice, including N=3 saline-treated controls, did not reach terminal disease endpoint because they died acutely following intraperitoneal luciferin injection for live animal imaging (see Discussion).

Despite these complications, ASO 1 prolonged all-cause mortality by 46% (median 234 vs. 160 dpi; Figure 5D, Table S2). Immediately after ICV injections, a sharp increase in BLI was observed in both saline- and ASO-treated mice, as a result of disease progression and/or inflammatory reaction to the surgical intervention (Figure 5D). BLI in mice treated with active ASO 1 declined to below the level in saline-treated animals at approximately three weeks post-dose, similar to time course at which NFL reversal was observed in the aforementioned experiment, albeit different ASOs were used (Figure 5A and 5C). Thereafter, BLI in saline-treated animals increased sharply up through terminal disease, while BLI in active ASO 1-treated animals remained low through terminal endpoint. In contrast to NfL, astrogliosis did not rebound at any timepoint after treatment, even as these mice developed typical prion disease on a similarly delayed schedule (medians 248 and 234 dpi in NfL and BLI experiments respectively, Figure 5B and 5D). These findings provide additional evidence that PrP-lowering can reverse pathological change.

### Chronic dosing initiated at different timepoints

Antiprion compounds with non-PrP-lowering mechanisms of action have been most effective in prion-infected mice when administered prophylactically or very early after prion infection, with diminished or no efficacy as animals approached symptoms (29, 32, 69, 70). In ASO experiments described above and previously (28), we intervened at various timepoints, but comparison of efficacy between timepoints is complicated because these experiments also differed in their number of doses and in their experimental endpoints (symptomatic versus terminal disease). We therefore designed a controlled experiment to assess how timing of intervention impacts the efficacy of PrP-lowering therapy. We also employed a chronic dosing paradigm, to more closely approximate clinical use of existing ASO therapies. A total of N=112 mice were infected with RML prions and groups of N=8 received doses of 500 μg active ASO 6 or saline every 90 days beginning at −7, 1, 28, 54, 78, 105, or 120 dpi. Across timepoints, all mice in this experiment were followed to a symptomatic endpoint by blinded raters. This contrasts with some of our prior experiments, in which late (83-129 dpi) treatment timepoints utilized a terminal endpoint (Figure 2, Figure 5, ref. (28)).

Based on our natural history study, the first four timepoints in this experiment (−7 to 54 dpi) precede rise in plasma NfL. 78 dpi falls between the 60 dpi timepoint where some animals show initial NfL rise, and 90 dpi where plasma NfL elevation is consistently evident in prion-infected animals. The latest timepoints, 105 and 120 dpi, occur after NfL pathology is clearly detectable and around the time when symptomatic changes can begin to be detected.

Across the first five timepoints, including 78 dpi, (Figure 6A-E, Table S4), we observed a dramatic increase in time to symptomatic endpoint, driven both by an increase in healthy lifespan as well as by a slowing of initial symptomatic decline, as reflected in weights, symptoms, and nest-building (Figure S6A-C). Survival did not differ significantly between these five early timepoint groups (*P* = 0.29, Type I ANOVA, Figure 6A-E), although weight loss and nest-building defects, but not observable symptoms, appeared to be delayed somewhat longer in the earliest-treated cohorts (Figure S6D-F). Initiation of treatment at later (105 and 120 dpi) timepoints, corresponding to 70% and 79% of the time to endpoint, still extended survival, although to a lesser extent compared to earlier timepoints (Figure 6F-G, Table S4). The effect size at 120 dpi observed in this experiment is smaller than our 120 dpi interventions against

RML prions in which animals were followed to a terminal disease endpoint (Figure 2D and ref. (28)), and more similar to our result for 123 dpi intervention against RML prions with a symptomatic endpoint (Table 3), suggesting that different endpoints explain the different outcomes between experiments at this timepoint. Overall, late (105-120 dpi) treatment increased survival by 19% (median 175 dpi vs. 147 dpi across all saline controls; Figure 6H). Meanwhile early (≤78 dpi) initiation, when paired with chronic treatment, was able to drive a striking survival increase of about 3.0x (median 437 vs. 147 dpi across all saline controls), on par with the benefit we observed with genetic reduction of PrP in RML-infected heterozygous knockout mice (Figure 6H).

### Intervention at the symptomatic disease stage

In our natural history study, we observed suggestive or nominally significant group-wise differences between RML prion-infected and uninfected animals in terms of observation scores, rotarod performance, and nest-building by 120 dpi (Figure 4). This timepoint may, however, still precede the development of obvious individual symptoms in many animals (Figure S4, S5). We therefore undertook a series of later treatments overlapping the frankly symptomatic phase of RML prion disease. A total of N=96 mice were infected with RML prions, and groups of N=12 received a single dose of 500 μg active ASO 6 or saline at 120, 132, 143, or 156 dpi, and were followed to a terminal disease endpoint by blinded raters. As for previous experiments with a terminal endpoint (Figures 2, 5, and ref. (28)), treatment at 120 dpi extended survival of a majority of animals (Figure 7A), allowed some recovery of lost weight (Figure 7B) and attenuated symptom accumulation and loss of nest-building (Figure 7C-D).

By the 132 and 143 dpi timepoints, corresponding to 81% and 85% of the time to terminal endpoint, most or all (22/23 and 23/23 surviving animals, respectively) had already declined from their individual peak weights. By the 143 dpi timepoint, nest-building defects were also evident (Figure 7). At these timepoints, ASO treatment was effective in only a minority of animals. 35% of ASO-treated animals survived the immediate post-surgical period, living >10% (~17 days) longer than saline-treated controls. Those that did so lived considerably longer (mean 85 days), albeit without any measurable recovery in body weight or nest building (Figure 7E-L). By 156 dpi, when 7/23 (30%) of mice intended for treatment had already reached the terminal disease endpoint, PrP-lowering therapy had no effect (Figure 7M-P, Table S5).

## Discussion

PrP lowering is a longstanding therapeutic hypothesis. We recently reported that PrP-lowering ASOs are effective against prion disease. The present results expand on and broaden our previous findings, outlining the parameters that govern the efficacy of PrP-lowering therapies in prion disease.

We confirmed that PrP-lowering ASOs extend survival of an intracerebrally inoculated model of prion disease in both prophylactic and delayed treatment paradigms, while non-PrP-targeting ASOs do not. These data replicate across ASO chemistries, confirm our previous findings (28), and build confidence that the efficacy of these molecules is RNAse H-mediated. These data provide important clarification on ASO mechanism of action in prion disease, as oligonucleotides have been shown to sequence-independently interact with prion protein *in vitro* (71), have been shown to reduce prion load *in cellulo* independent of impact on PrP levels (72, 73), and have been shown to modulate prion disease in animals when pre-incubated with prion inoculum prior to injection, or when used to treat a prion disease in peripheral tissues (72). Our own studies have confirmed high affinity *in vitro* interactions between PrP and the same ASOs used in our *in vivo* studies, across chemistries and regardless of sequence (58). Given the hazards surrounding interpretation of oligonucleotide experiments generally (74), and this background for PrP in particular, the confirmation here of an RNAse H-mediated, rather than aptameric, mechanism of action *in vivo* is important.

Indeed, the PrP-lowering mechanism of action of ASOs has specific implications for advancement of oligonucleotide therapeutics for prion disease. Our data show that ASO treatment was effective against five out of five prion strains tested, suggesting a universality of PrP lowering across subtypes of prion disease that is yet to be established for any other therapeutic strategy. Our data further suggest that degree of reduction of prion protein RNA correlates with efficacy against prion disease *in vivo*. In dose-response studies, we observed a clear relationship between degree of *Prnp* RNA suppression and extension of survival. Our experiments identified no minimum threshold to effect, with a clear survival benefit from even transient 21% knockdown, consistent with the continuous dose-response relationship postulated from genetic models (8). Both pharmacologic and genetic reduction of PrP were substantially effective against five of five prion strains tested, and we did not observe emergence of drug resistance. The observation of a quantitative relationship between *Prnp* RNA knockdown and efficacy reinforces potency of PrP reduction as a key criterion for discovery and prioritization of oligonucleotide therapeutics for prion disease. These data will also be important to the qualification of CSF PrP concentration as a pharmacodynamic biomarker for PrP-lowering drugs (44, 75, 76). The validation of ASOs’ mechanism of action *in vivo*, the tight relationship between degree of PrP lowering and disease delay, and the efficacy across prion strains, observed here all support the disease relevance of this biomarker (2).

The efficacy of previous antiprion therapies has depended critically upon the timepoint when treatment was initiated (29, 32, 69, 70). To better define disease timepoints in our animal model, we conducted natural history and biomarker studies in intracerebrally RML prion-inoculated mice. Biomarker evidence of pathology became clear well before the onset of detectable symptoms. Astrocytosis was detected by bioluminescence imaging beginning at 73-81 dpi, while plasma NfL became elevated in some animals by 60 and in all animals by 90 dpi. Notably, the disease-associated rise in both biomarkers could be measurably reversed by a single ASO treatment. In contrast, rotarod impairment became nominally detectable at 120 dpi, observable symptom profiles emerged by 116-135 dpi, and weight loss did not become obvious until 148 dpi. This is consistent with previous reports indicating neuroinflammatory changes can be observed by ~55-60 dpi (67, 68, 77), neuronal damage between 60-75 dpi (78), and behavioral or motor changes by ~105 dpi or later (77, 79).

Overall, PrP-lowering therapy showed efficacy across a wide range of treatment timepoints. Chronic dosing initiated at pre-symptomatic timepoints up to early detectable pathology (≤78 dpi) tripled the time to a symptomatic endpoint (an increase of 290 days), both by extending healthy life and slowing initial decline. This matches the effect observed here and elsewhere (5–7) in heterozygous PrP knockout mice, and is consistent with PrP expression being required for both prion propagation and neurotoxicity (80, 81). Intervention at neuropathological timepoints approaching the time of earliest detectable symptomatic changes (83-120 dpi) also increased survival time, with reversal of neuronal damage and astrocytosis markers and some recovery of initial weight loss. At these pathological timepoints, we observed a modest delay (~1 month) in time to a symptomatic endpoint (accumulation of five prion disease symptoms), and a more profound delay (~3 months) in time to terminal disease (with criteria including 15-20% body weight loss).

At frankly symptomatic timepoints (132-143 dpi), we observed a ~85 day delay in terminal disease in approximately one third of animals (8/23), without reversal of weight loss, nesting, or symptomatic changes. At the most advanced symptomatic endpoint (156 dpi), no benefit was observed. Both the bimodal outcomes observed at 132 and 143 dpi, and the lack of effect seen at 156 dpi, raise the possibility that treatment at these stages of disease may not allow not enough time for ASOs to take effect prior to most or all animals reaching endpoint. Although ASOs engage RNA targets within one week (Figure 1 and ref. (82)), our biomarker studies suggest a three-week lag time for this target engagement to impact established pathology (Figure 5). While the half-lives of PrP and misfolded prions are reported to be on the order of 1-5 days (10, 83), recovery from prion neurotoxicity may be more gradual. Broadly, the spectrum of outcomes observed at different timepoints may reflect accumulation of irreversible damage during the disease course, and may suggest the value of testing more aggressive dosing regimens when treatment is initiated later in the disease course.

Consistent with previous reports (22, 28), not all preclinical ASOs were tolerated by mice with established prion neuropathology. These animals experienced an acute, accelerated decline following surgery that was phenotypically distinct from prion disease, as we and others have reported previously (22, 28). This phenomenon may reflect the limited screening and optimization undertaken to identify these tool compounds. Studies to elucidate the mechanism at work are ongoing. We also observed that animals with advanced prion disease often died immediately after luciferin injection for live animal imaging. Such deaths have not been reported before in Tg(*Gfa*p-luc) mice (46), have not been observed during our extensive experience of BLI studies in non-prion animals, and were never observed in our uninfected controls. Three saline-treated animals succumbed in this manner, ruling out a specific interaction between ASOs and luciferin, but instead suggesting the fragility of mice with advanced prion infection to experimental manipulation.

Our study has important limitations. While we investigated two biomarkers and a large battery of symptom endpoints, our understanding of the natural history of experimental prion disease is by no means exhaustive, and other approaches have nominated putative pathological and symptomatic changes somewhat earlier than we observed here (67, 77, 79). While we consistently observed an overall survival benefit to PrP-lowering therapy across nearly all paradigms tested, sometimes only a subset of mice benefitted, and the magnitude of therapeutic benefit observed sometimes varied between nearly identical experiments. This could reflect many contributing factors including variability in ICV dosing efficiency, human error in animal evaluation, and the imperfect tolerability of the ASO tool compounds employed.

In the reported studies, we rely on *Prnp* RNA reduction as a proxy for protein reduction. Appropriateness of this proxy is supported by our previous data characterizing ASO-mediated reduction in both RNA and protein with a subset of these same tool compounds (28), and by historical data showing that in *Prnp^+/−^* mice, 50% wild-type *Prnp* RNA levels correspond to 50% protein levels (8). Lack of efficacy across non-targeting ASOs, and the close tracking of ASO-mediated survival benefit with heterozygous *Prnp* knockout, further build confidence that our results are driven by substrate reduction, rather than an orthogonal mechanism of action.

We chose to use wild-type mice intracerebrally inoculated with prions for the majority of our studies. A number of factors made this model more appropriate than available alternatives. Prion-inoculated wild-type animals follow a well-established, phenotypically relevant prion disease course culminating in terminal illness. They propagate transmissible prions, and their brains develop characteristic histopathological and biochemical hallmarks that together unambiguously signify prion disease (57). Importantly, these phenotypic and molecular features develop in the context of endogenous PrP expression levels, which may have special relevance when assessing a PrP-lowering treatment: genetic data suggest that the therapeutic benefit of reducing PrP levels by half is likely to be smaller in the context of overexpression than in the context of endogenous expression (8). By contrast, mouse models that develop spontaneous prion disease generally have one or more of the following limitations: overexpression of PrP, development of only subtle disease signs, highly variable times to disease, and/or lack of transmissible prions (57). In addition, use of wild-type animals is appropriate to the mechanism of action of ASOs. Because ASOs are active in the nucleus (84), some active ASO compounds target intronic sequences, which may not be present in transgenic models (85). Even a transgene containing the targeted sequence, depending on its construction, may produce a truncated RNA. This altered RNA, in turn, may be less potently targeted than the wild-type RNA by ASOs initially screened for potency in wild-type mice. To counter our reliance on inoculated wild-type animals, we have sought to capture a diversity of experimental paradigms, including a variety of strains, timepoints, doses and endpoints, to build confidence in our results.

The effectiveness of a given PrP-lowering dosing regimen may vary depending on the stage of the disease, suggesting that dose regimens and trial endpoints may need to be adjusted depending on the clinical profile of the trial population. Nevertheless, our findings provide basis for optimism that PrP lowering may be a promising therapeutic strategy, both for prophylaxis against prion disease onset in at-risk individuals with no evidence of disease process underway (43, 44), and for treatment of active prion disease, during either prodromal or manifest disease.

## Author contributions

Conceived and designed the studies: SMV, DEC, HBK, HTZ, EVM, JBC, GAC

Analyzed the data: EVM, SMV, HTZ, DEC, JBK

Provided key methods or reagents: GAC, JK, JBK, JYM, HW, DM, JA, KDH

Performed the experiments: SMV, DEC, HTZ, EVM, JL, JOM, RP, SG

Supervised the research: SMV, DEC, HBK, MPK, TC, JS, MB, SLS, ROK

Wrote the manuscript: SMV, EVM, HTZ, HBK, DEC

Reviewed, edited and approved the final manuscript: All authors

## Acknowledgements

We thank Rob Pulido, Cecilia Monteiro, Anne Smith, and Tiffany Baumann for critical review of the manuscript, Karli Ikeda-Lee and Amanda Sevilleja for technical assistance, Johnnatan Tamayo, Samantha Allhands, and Donna Sipe for vivarium assistance.

## SUPPLEMENTARY MATERIALS

**Figure S1.**
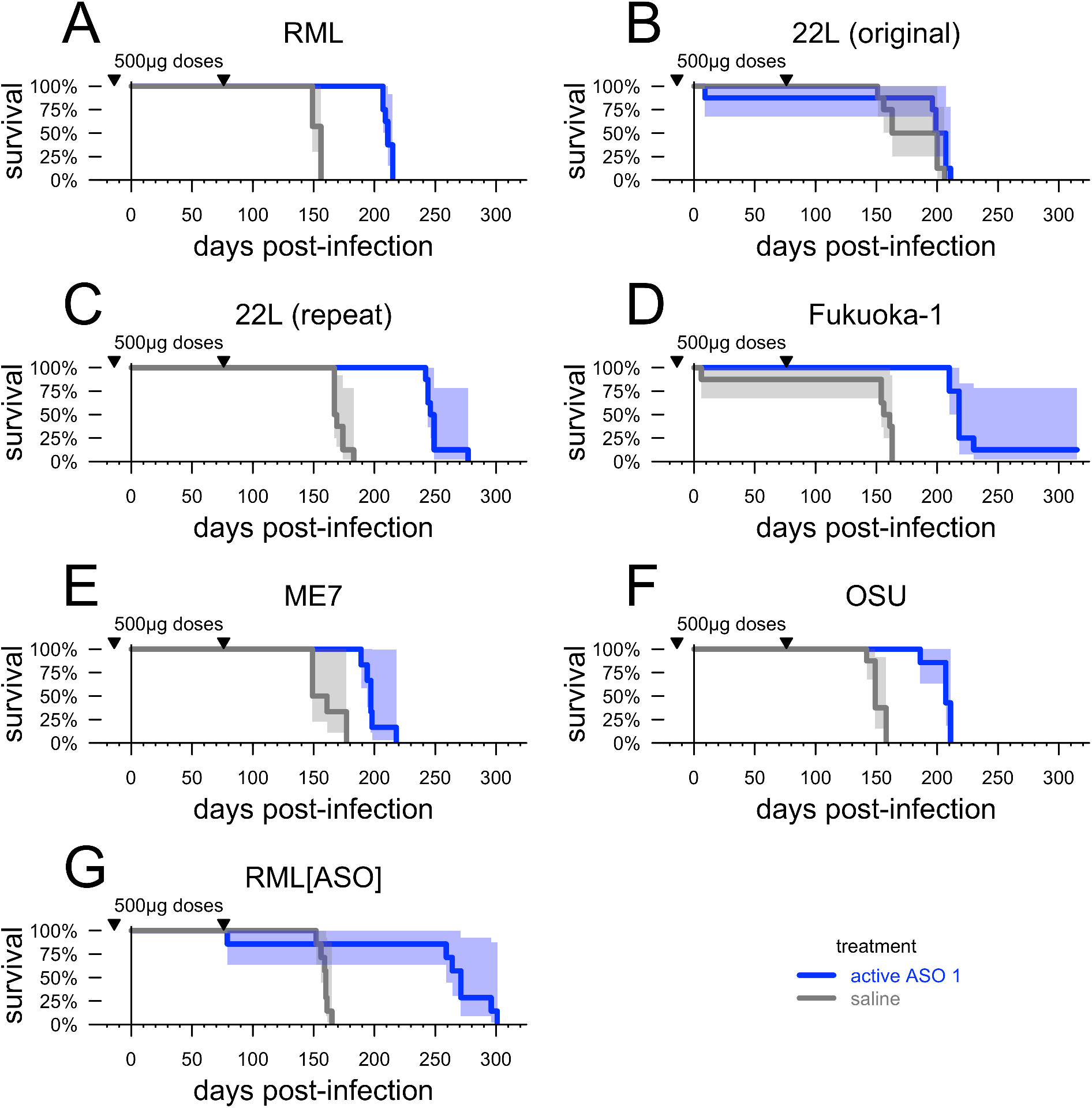
Survival curves for prophylactically ASO-treated animals across prion strains. Data summarized in Table 2. Shaded areas represent 95% confidence intervals. In the original 22L experiment (B), we observed only a marginal increase in survival time in ASO-treated animals (+12%, P = 0.09, two-sided log-rank test). We suspected an experimental error because the distribution of survival times was bimodal among control animals: 4/4 saline-treated animals in one cage succumbed at 158±6 dpi, while 4/4 saline-treated animals in the other cage succumbed at 202±3 dpi. This latter cage had been labeled “E” while an adjacent active ASO-treated cage had been labeled “F”, leading us to suspect the two cage cards had been swapped and that some of the 22L control animals had in fact received one dose of ASO. We repeated the experiment, again with blinded veterinary technicians performing all animal evaluations, and obtained the result in panel (C), which is summarized in Table 2.

**Figure S2.**
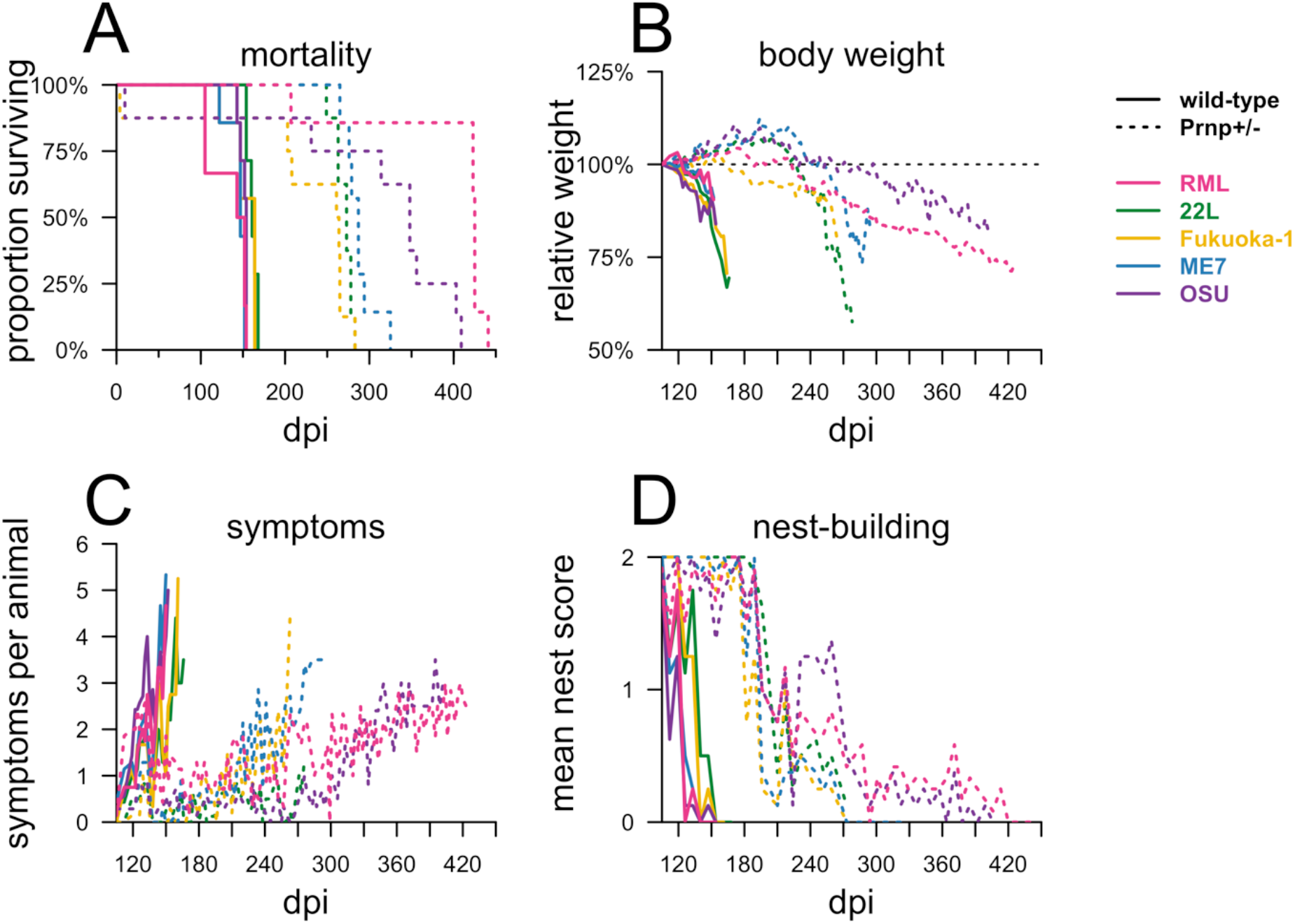
Endpoints in heterozygous PrP knockout mice infected with five prion strains. Details on animals summarized in the right half of Table 2. **A)** survival, **B)** body weights relative to 105 dpi baselines (relative rather than absolute weights are used because the cohorts contain different proportions of male and female mice), **C)** mean symptom count per animal, and **D)** mean nest score.

**Figure S3.**
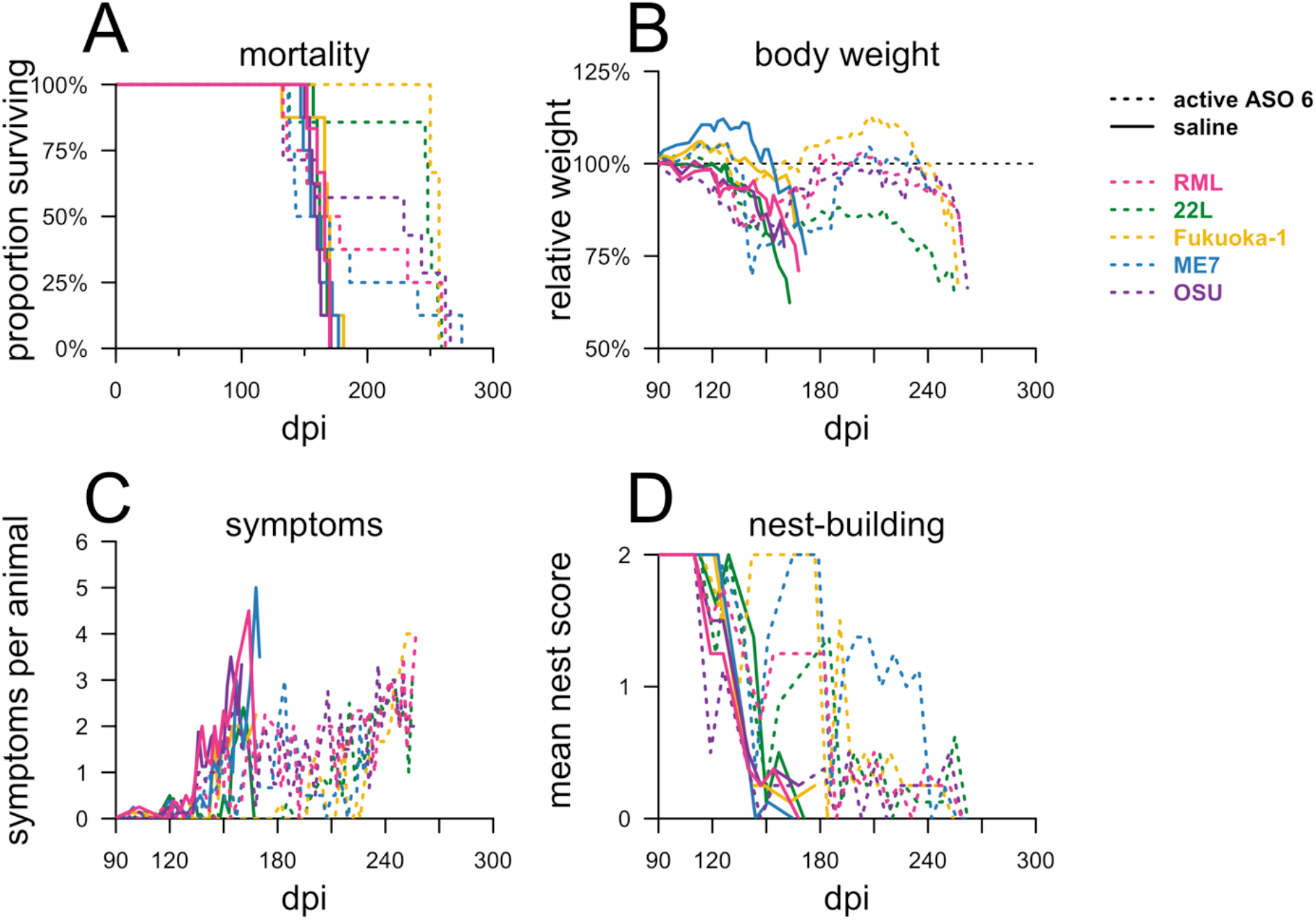
Endpoints in mice infected with five prion strains receiving late ASO or saline treatment. Details on animals summarized in Table 3. **A)** survival, **B)** body weights relative to 105 dpi baselines, **C)** mean symptom count per animal, and **D)** mean nest score.

**Figure S4.**
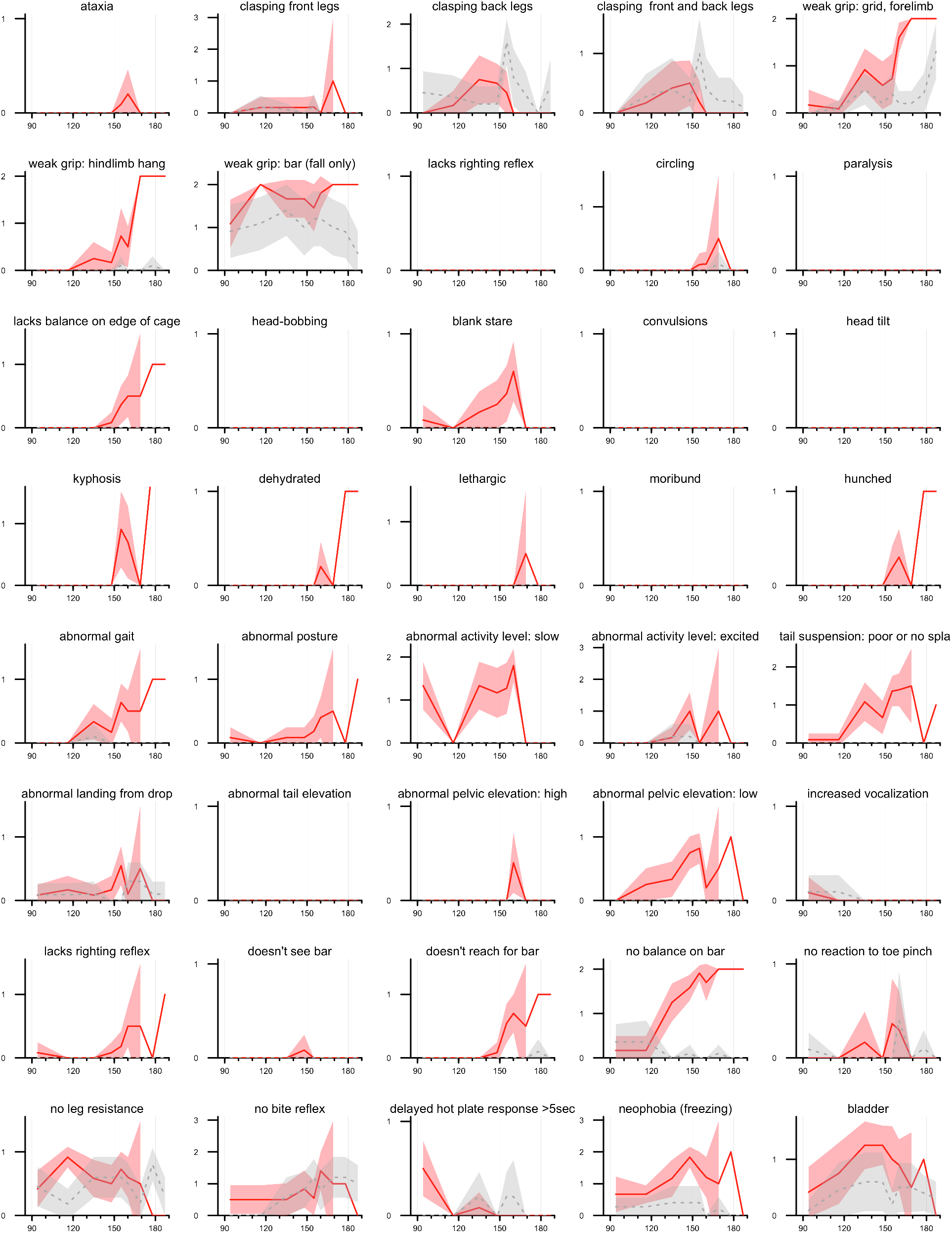
Behavioral observations in the natural history of RML prion infection. Data from Figure 4C broken into all N=40 Individual behavioral observations (listed in Table S2).

**Figure S5.**
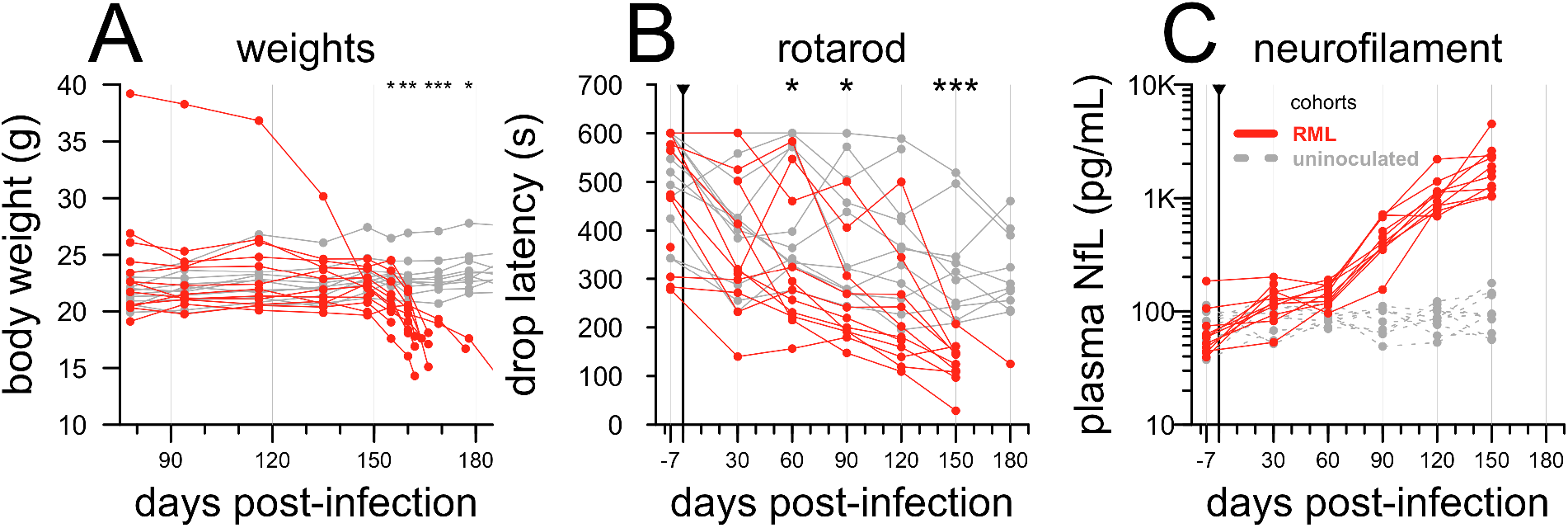
Raw individual natural history endpoints. Alternative visualizations of data from Figure 4: **A)** individual body weights, **B)** individual mean rotarod latency, and **C)** individual plasma NfL concentrations (note log y axis).

**Figure S6.**
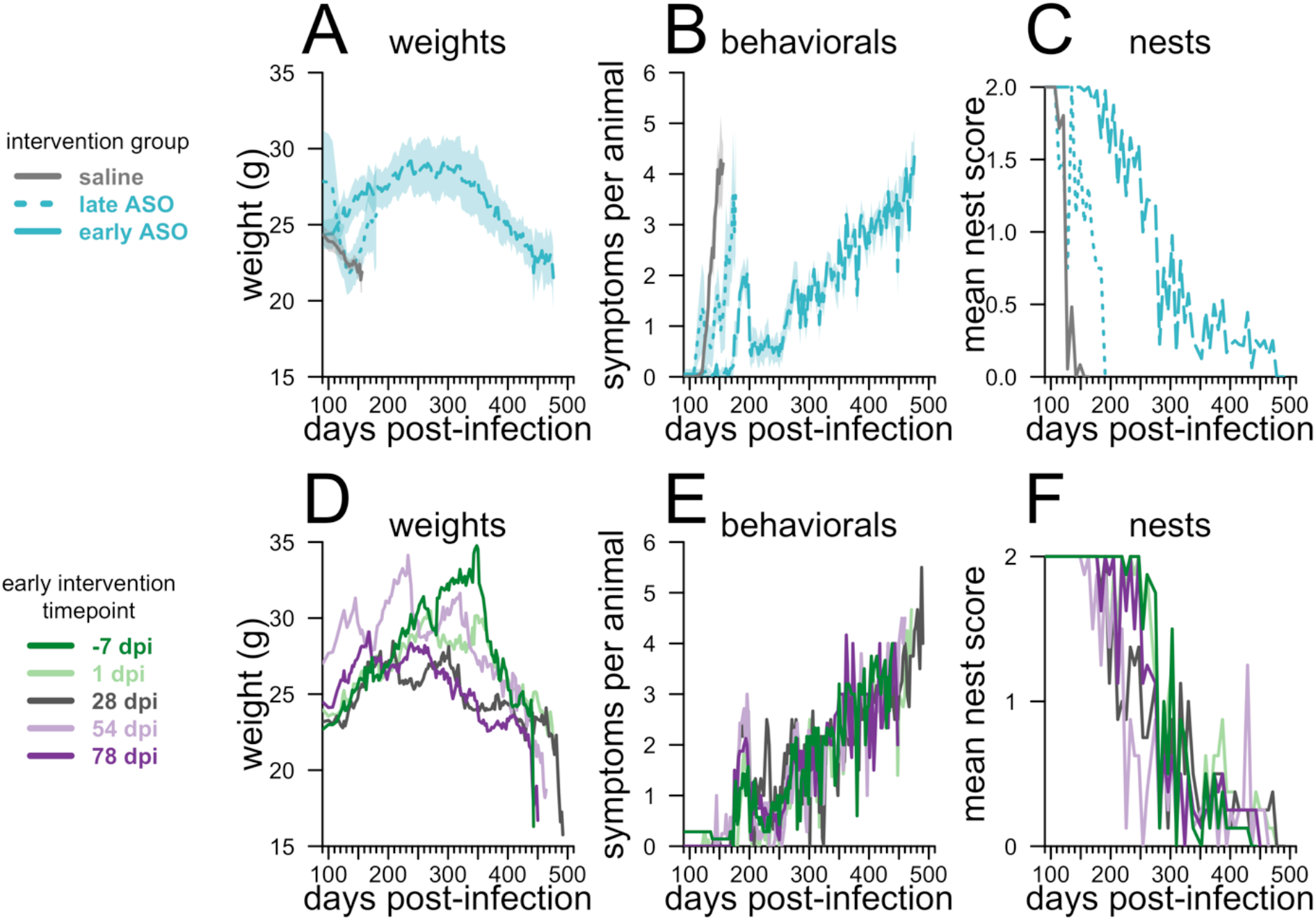
Endpoints in mice receiving chronic ASO or saline treatment beginning at different timepoints. Details on animals summarized in Figure 6: weights (**A,D**), mean symptom count per animal (**B,E**), and nest score (**C,F**). **A-C**) animals are grouped by saline or early (−7 to 78 dpi) versus late (105 and 120 dpi) ASO treatment initiation, as in Figure 6B. **D-F**) early intervention (−7 to 78 dpi) animals are grouped by individual intervention timepoint.

**Table S1.** Detailed criteria and performance statistics for behavioral tests. Building on commonly used neurological evaluations in the prion field^2^, we initially (experiments reported in Raymond et al^3^ and in Figure 2D-F) evaluated animals for six neurological signs: tremor, ataxia, difficulty righting, tail rigidity, blank stare, and hindlimb weakness. Based on those initial experiences, after removing signs found non-contributory, grouping those found redundant, and adding signs sometimes noted in comments, our behavioral battery was revised to the above list for all other experiments at the Broad Institute reported herein (Figure 2A-C, 3, 6-7, and Tables 2-3). Raters were asked to assess each animal individually for 15 seconds on a cage lid and to record only observations made during this period. Each sign was then recorded as a 1 or 0 in a blank set of spreadsheet columns. Specific criteria or guidance were provided to raters for a subset of tests and are displayed above; other tests were never described in greater detail verbally but were conveyed through hands-on training. In order to assess the baseline performance characteristics of each test, we retrospectively analyzed N=119 saline-treated, RML prion-infected control animals across all experiments. For each animal, time of onset of each symptom was defined as the first timepoint where the symptom was i) observed at that timepoint, and ii) observed in at least half of subsequent observations of that animal. Displayed are the mean±sd of the lead time (time from onset of specific symptom to prion disease endpoint) and the proportion of animals in which the symptom was ever observed.

| behavioral test | criteria/guidance | lead time to disease endpoint (mean±sd) | proportion of animals ever showing symptom |
| --- | --- | --- | --- |
| tremor | continuous tremor visible to the naked eye (periodic tremors, or tremors that can be felt but not seen, do not count) | 20.1 ± 10.2 | 98% |
| reduced activity | moves <50% as much or <50% as fast as a normal mouse | 18.8 ± 8.3 | 97% |
| scruff/poor grooming | full body scruff (neck only does not count); mice with boosts cannot be scored | 8.7 ± 6.7 | 86% |
| irregular gait/hindlimb weakness | — | 8.0 ± 5.1 | 73% |
| blank stare | — | 4.3 ± 3.1 | 55% |
| hunched posture | — | 4.0 ± 3.8 | 40% |
| poor body condition | body condition score<sup>1</sup> ≤2 | 2.7 ± 1.5 | 5% |
| difficulty righting | perturb the mouse by lying on its side and see if it stands up again | 2.6 ± 1.5 | 8% |

**Table S2.** Endpoint data for animals in Figures 2-5 in tabular form. Only animals that reached pre-specified euthanasia endpoint are included here. For overall mortality data see survival curves in the main figures or the online data repository described in Methods. *In the uninoculated group, no animals developed disease; the experiment was terminated at 190 dpi.

| Experiment | cohort | mean±sd | N | $\Delta$ |
| --- | --- | --- | --- | --- |
| MOE ASO prophylactic (Figure 2A-C) | saline | 150.7 ± 5.5 | 9 | — |
| | control ASO 4 | 157.9 ± 4.9 | 10 | +5% |
| | active ASO 5 | 304.8 ± 19.9 | 6 | +102% |
| | active ASO 6 | 276.8 ± 11.6 | 9 | +84% |
| MOE ASO 120 dpi (Figure 2D-F) | saline | 161.7 ± 7.0 | 9 | — |
| | control ASO 4 | 160.2 ± 11.0 | 9 | -1% |
| | active ASO 5 | 139.5 ± 5.3 | 4 | -14% |
| | active ASO 6 | 275.2 ± 3.2 | 8 | +70% |
| Dose response (Figure 3) | saline | 152.0 ± 5.3 | 8 | — |
| | 30 μg | 173.6 ± 7.7 | 7 | +14% |
| | 100 μg | 194.0 ± 6.7 | 6 | +28% |
| | 300 μg | 207.2 ± 4.3 | 6 | +36% |
| | 500 μg | 226.4 ± 14.7 | 7 | +49% |
| Natural history (Figure 4) | uninoculated | 190* | 0 | — |
| | RML | 164.8 ± 9.6 | 12 | — |
| NfL (Figure 5A-B) | saline | 160.7 ± 9.9 | 10 | 0% |
| | active ASO 6 | 253.0 ± 11.7 | 8 | 57% |
| BLI (Figure 5C-D) | saline | 156.2 ± 12.1 | 8 | 0% |
| | control ASO 3 | 155.7 ± 9.8 | 3 | 0% |
|  | active ASO 2 | 189 | 1 | +21% |
| | active ASO 1 | 234.7 ± 18.1 | 7 | +50% |

**Table S3.** Measures observed in the natural history study.

| category | observations | scoring |
| --- | --- | --- |
| neurological | ataxia | 1: difficulty getting across cage and /or falling over |
| neurological | clasping front legs | 1: almost 2: full clasp |
| neurological | clasping back legs | 1: almost 2: full clasp |
| neurological | clasping front and back legs | 1: almost 2: full clasp |
| neurological | weak grip: grid, forelimb | 1: weak or moderate 2: absent |
| neurological | weak grip: hindlimb hang | 1: moderate (fall 1 out of 2 times) 2: weak fall 2 out of 2 tries (2x) |
| neurological | weak grip: bar (fall only) | 1: moderate (fall 1 out of 2 times) 2: weak fall 2 out of 2 tries (2x) |
| neurological | lacks righting reflex (flipped on back by tail, not contact right reflex) | 1: poor: all but head will roll over when tail is twisted. 2: lacking: can't right |
| neurological | circling | 1: present |
| neurological | paralysis | 1: paresis (limited range of motion for walking only, curled paw may be present, one limb non-weight bearing) 2: paralysis (cannot move) |
| neurological | lacks balance on edge of cage | 1: present |
| neurological | head-bobbing | 1: present |
| neurological | blank stare | 1: present at initial box evaluation |
| neurological | convulsions | 1: present |
| neurological | head tilt | 1: present |
| neurological | kyphosis | 1: slight to moderate 2: severe |
| non-neurological | dehydrated | 1: present when nape pinched and/or sunken eyes |
| non-neurological | lethargic | 1: present (immobile even after poked by finger) |
| non-neurological | moribund | 1: present (near death) |
| non-neurological | hunched | 1: present, not kyphosis, front and hind feet are close together when standing, not applicable to resting posture |
| SHIRPA | abnormal gait | 1: wobble due to wide hind limb stance |
| SHIRPA | abnormal posture | 1: present |
| SHIRPA | abnormal activity level: slow | 1: present in home box only 2: no exploring in test box |
| SHIRPA | abnormal activity level: excited | 1: moderate (noticed in home box or behavior box) 2: severe, may include jumping |
| SHIRPA | tail suspension: poor or no | 1: poor 2: none or abnormally wide |
|  | splay |  |
| SHIRPA | abnormal landing from drop | 1: falls on tail or back from bar only (not drop from tail suspension) |
| SHIRPA | abnormal tail elevation | 1: present |
| SHIRPA | abnormal pelvic elevation: high | 1: present |
| SHIRPA | abnormal pelvic elevation: low | 1: present |
| SHIRPA | increased vocalization | 1: present |
| SHIRPA | lacks righting reflex | 1: lacking |
| SHIRPA | doesn't see bar | 1: present |
| SHIRPA | doesn't reach for bar | 1: present |
| SHIRPA | no balance on bar | 1: moderate: can balance after multiple tries or can only balance a second or two. 2: cannot balance |
| SHIRPA | no reaction to toe pinch | 1: flinch 2: no reaction (zero: pulls paw away immediately) |
| SHIRPA | no leg resistance | 1: doesn't react to pressure against paws (zero if any paw pushes back) |
| SHIRPA | no bite reflex | 1: poor: notices but doesn't bite 2: no reaction (normal is immediate bite) |
| SHIRPA | delayed hot plate response >5sec | 1: longer than 5 seconds |
| SHIRPA | neophobia (freezing) | 1: any episode of freezing in test box (vs home box) 2: paralyzing fear |
| SHIRPA | bladder | 0: not palpable (np) or good abdominal tone (gt), 1: small or medium 2: large or huge. |

**Table S4.** Endpoint times by treatment timepoint in chronic dosing study. Data from Figure 6 presented in tabular format. In contrast to the Results text, this presentation of the data uses mean±sd as opposed to median and includes only animals reaching pre-specified euthanasia endpoint. For overall mortality data see survival curves in the main figures or the online data repository described in Methods.

| timepoint | saline | N | active ASO 6 | N | $\Delta$ |
| --- | --- | --- | --- | --- | --- |
| | survival dpi (mean±sd) | | survival dpi (mean±sd) | | |
| -7 | 143±4 | 7 | 394±77 | 4 | +175% |
| 1 | 145±3 | 8 | 462±18 | 6 | +219% |
| 28 | 145±11 | 6 | 426±109 | 7 | +194% |
| 54 | 150±5 | 8 | 377±124 | 5 | +151% |
| 78 | 149±7 | 7 | 406±47 | 4 | +172% |
| 105 | 151±4 | 8 | 184±5 | 5 | +21% |
| 120 | 152±7 | 8 | 172±8 | 7 | +13% |

**Table S5.** Survival times by treatment timepoint in symptomatic intervention study. Data from Figure 7 presented in tabular format. In contrast to the Results text, this presentation of the data uses mean±sd as opposed to median and includes only animals reaching pre-specified euthanasia endpoint. For overall mortality data see survival curves in the main figures or the online data repository described in Methods.

| timepoint | saline | N | active ASO 6 | N | $\Delta$ |
| --- | --- | --- | --- | --- | --- |
| | survival dpi (mean±sd) | | survival dpi (mean±sd) | | |
| 120 | 167±9 | 12 | 243±50 | 7 | +46% |
| 132 | 164±14 | 8 | 187±57 | 8 | +14% |
| 143 | 168±7 | 12 | 207±65 | 7 | +23% |
| 156 | 165±5 | 5 | 163±5 | 7 | -1% |

